# UBA52 is crucial in HSP90 ubiquitylation and neurodegenerative signaling during early phase of Parkinson’s disease

**DOI:** 10.1101/2022.08.17.504224

**Authors:** Shubhangini Tiwari, Abhishek Singh, Parul Gupta, Sarika Singh

**Affiliations:** Division of Neuroscience and Ageing Biology, Division of Toxicology and Experimental Medicine, CSIR-Central Drug Research Institute, Lucknow 226031, India; Academy of Scientific & Innovative Research (AcSIR), Ghaziabad 201002, India

**Keywords:** Parkinson’s Disease (PD), Protein Aggregation, Endoplasmic Reticulum Stress, Ubiquitin-Proteasome System (UPS), Ubiquitin-60S Ribosomal Protein L40 (UBA52), Dopaminergic Neuronal Death

## Abstract

Protein aggregation is one of the major pathological events in age-related Parkinson’s disease (PD) pathology, predominantly regulated by the ubiquitin-proteasome system (UPS). UPS essentially requires core component ubiquitin however, its role in PD pathology is obscure. This study aimed to investigate the role of ubiquitin encoding genes in the early phase of PD pathology. Wild-type human Myc-α-synuclein transfected neurons, α-synuclein-PFFs treated cells, rotenone-induced sporadic models of PD and SNCA C57BL/6J-Tg (Th-SNCA*A30P*A53T)39 Eric/J transgenic mice showed downregulated level of UBA52 in conjunction with significant downregulation of tyrosine hydroxylase (TH) and neuronal death. *In silico* predictions, mass spectrometric analysis and co-immunoprecipitation findings suggested strong interaction of UBA52 with α-synuclein, HSP90 and E3-ubiquitin ligase CHIP, besides its co-localization with α-synuclein in the mitochondrion. Next, *in vitro* ubiquitylation assay indicated an imperative requirement of the lysine-63 residue of UBA52 in CHIP-mediated HSP90 ubiquitylation. Myc-UBA52 expressed neurons exhibited the downregulated α-synuclein protein abundance with increased TH and restored proteasome activity during the diseased condition. Furthermore, Myc-UBA52 expression inhibited the augmented HSP90 protein level along with its various client proteins, HSP75 (homologue of HSP90 in mitochondrion) and ER stress-related markers during early PD. Taken together, data highlights the critical role of UBA52 in HSP90 ubiquitylation in parallel to its potential contribution to the modulation of various disease-related neurodegenerative signaling targets during the early phase of PD pathology.

## Introduction

Age-related movement disorder Parkinson’s disease (PD) is caused due to the loss in dopaminergic neurons of substantia nigra in the midbrain (Lang & Lozano 1998; Tiwari & Singh 2020; Singh & Dikshit 2007). The neuropathological hallmarks of the disease include Lewy bodies containing protein (α-synuclein, ubiquitin, chaperones) aggregates, reflecting the impaired protein degradation mechanisms like ubiquitin-proteasome system (UPS) and autophagy (Tiwari & Singh 2020). Pertained organelle endoplasmic reticulum (ER) dysfunction has been reported in PD by us and others (Goswami et al. 2016; Costa et al. 2020; Kovaleva & Saarma 2021; Holtz & O’Malley 2003; Greenamyre & Hastings 2004; Ryu et al. 2002; Cimato et al. 1997) that eventually leads to the dopaminergic neuronal death. In concurrence, recently we have shown the cardinal role of eukaryotic initiation factor 2α during the early phase of PD pathology (Gupta, Mishra, et al. 2021). The dysfunctional ER physiology leads to the accumulation of aberrant and unassembled proteins that need to be degraded by proteasome machinery / UPS. Reduction in proteasome activity during PD has also further indicated the critical role of UPS in disease pathogenesis (Singh et al. 2021; Bi et al. 2021). UPS assertively requires the attachment of ubiquitin moiety to the target protein for their degradation with the mandatory requirement of enzymes E1 (ubiquitin-activating enzyme), E2 (ubiquitin conjugation protein) and E3 (ubiquitin ligase), a process called ubiquitylation. Tagging of ubiquitin on the substrate proteins, through lysine residues, either leads to their degradation through proteasome or asserts their participation in various intracellular pathways (Tiwari & Singh 2020). Since ubiquitin moiety is an indispensable requirement of UPS, this study is focused on ubiquitin encoding genes during the early phase of PD pathology. Ubiquitin is a 76 amino acid-containing moiety encoded by the four genes-UBB, UBC, UBA52 and RPS27a. UBB and UBC encode for the polyubiquitin chains, whereas UBA52 and RPS27a are fusion proteins with ubiquitin residue on the N-terminus and ribosomal protein (L40 and S27a) on the C-terminus. In addition, the key participation of the related regulatory ubiquitin ligases has also been evaluated.

The role of UBA52 has been reported in other pathologies however, in PD pathology we for the first time report its critical role. In traumatic brain injury, altered mRNA and protein levels of UBA52 have been observed (Yao et al. 2007), while in diabetic nephropathy and hepatoma cell apoptosis, upregulated UBA52 was found (Kobayashi et al. 2016). In embryonic development also the role of UBA52 has been reported as UBA52 deficient mice exhibited decrease in protein synthesis, cell cycle arrest and death (Kobayashi et al. 2016).

The RPL40 subunit of UBA52 participates in the initiation, translation and elongation process through its interaction with the eukaryotic elongation factor-2 [eEF2] (Fernández-Pevida et al. 2012). Emerging shreds of evidence suggest that in PD, the phosphorylation of eEF2 takes place which promotes its dissociation from the ribosome and stalls the mRNA translation process, thereby stimulating the α-synuclein aggregation and consequent dopaminergic cytotoxicity (Jan et al. 2018). Although unreported, UBA52 and upregulated phosphorylated level of eEF2 might have neurodegenerative implications during PD pathology that needs to be studied and henceforth, obligated us to first understand the role of ubiquitin genes in PD pathology. Along with this prime objective, further, the study was extended to understand the implication of UBA52 in chaperones functioning and biochemical alterations, specifically related to the mitochondrion and ER organelle during PD pathology.

ER organelle is primarily responsible for folding of the nascent polypeptide with the help of chaperones and targets the misfolded proteins to UPS. Since we have already depicted the insinuation of ER signaling in PD, here in this study we evaluated the role of the involved chaperone in regulating protein machinery. HSP90 is one of the known predominant chaperones that co-localize with α-synuclein in the Lewy bodies during α-synucleinopathies and is associated with impaired proteasome function (Uryu et al. 2006). The activity of HSP90 is also modified by the co-chaperone HSC70. (Burmann et al. 2020) previously reported that the selective inhibition of α-synuclein interaction with HSC70 as well as with HSP90, especially with HSP90β, leads to its re-localization in the mitochondrion and subsequent formation of protein aggregates. It has been reported that during cellular stress, such aberrant and misfolded proteins are recognized by molecular chaperones which assist the E3 ligases such as the C-terminal of Hsp70 interacting protein (CHIP), Parkin, Siah1/2 and others in their ubiquitylation and degradation by the UPS (Friesen et al. 2017). E3 ligase, CHIP interacts with chaperones HSP70 & HSP90 to participate in the protein quality control (PQC) and UPS-mediated degradation of HSP90 & α-synuclein, a component of Lewy body (Zhang et al. 2005). In addition to CHIP, another E3 ligase Parkin also interacts with and ubiquitylates α-synuclein-interacting protein, synphilin-1. The co-expression of synphilin-1, α-synuclein and Parkin leads to the formation of ubiquitin-positive Lewy body inclusions, suggesting a key role of Parkin in ubiquitylating proteins and participation in Lewy body formation (Chung et al. 2001). Additionally, it has also been suggested that PINK1 phosphorylates both ubiquitin and Parkin to fully activate Parkin for its utmost E3 ligase activity, leading to its translocation to the damaged mitochondrion in order to regulate the pathogenic events (Koyano et al. 2014), suggesting the close association of ubiquitin and E3 ligases in energy-dependent degenerative mechanisms.

In view of the conjectural role of UBA52 in neuronal viability, its interference in the protein translation and PQC through chaperones, the present study has been focused to gain insight into the versatile role of UBA52 during early the phase of sporadic PD, utilising both the cellular and experimental rat models.

## Materials and Methods

### Chemicals and Antibodies

Please refer to Suppl. Table-1.

### Cell Culture, differentiation and treatments

The human neuroblastoma SH-SY5Y, rat neuroblastoma N2a and rat pheochromocytoma PC12 cells were maintained in Dulbecco’s modified Eagle’s medium (DMEM) and F12 (1:1) media supplemented with penicillin/streptomycin and 10% fetal bovine serum (FBS) or 10% FBS and 5% horse serum for PC12 cells under the atmosphere of 37°C and 5% CO_2_. Cells were cultured in culture flasks or culture well plates at 37°C in a water-saturated atmosphere of 95% air and 5% CO_2_. Neurotoxin (rotenone) treatment was given in SH-SY5Y for 24 h to mimic the diseased conditions (Tanner et al. 2011; Xicoy et al. 2017). For α-synuclein-PFFs (preformed fibrils)-induced experiments, 5μg/ml of recombinant human α-synuclein protein PFF (Abcam, ab218819) was given in SH-SY5Y cells and then processed for immunofluorescence labelling and co-localization studies (Ross et al. 2020). The basal proteasome activity of SH-SY5Y cells was inhibited with MG132 (cell-permeable proteasome inhibitor) treatment at a concentration of 5μM for 6 h, prior to any other respective experimental set-up/treatment. In a set of experiments, the cells were treated with tunicamycin (ER stress inducer) at 1μM for 24 h, as a positive control for ER stress. Experiments were also conducted in differentiated SH-SY5Y cells, N2a cells and PC12 cells, cultured in DMEM and F12 (1:1) media supplemented with penicillin/streptomycin and 1% FBS at 37°C and 5% CO_2_. *All-trans* retinoic acid (ATRA, 10μM) was added to the differentiating media of SH-SY5Y and N2a cells every alternate day for 7 days before assessing for morphological changes in differentiated cells (Cheung et al. 2009; S Narasimhan et al. 2020), followed by rotenone treatment to SH-SY5Y (500 nM) and N2a (1μM) for 24 h. For PC12 cells, 50ng/ml nerve growth factor (NGF) was added every alternate day for 7 days to induce differentiation and rotenone treatment (1μM) was given for 24 h (Cimato et al. 1997). For chronic *in vitro* studies in SH-SY5Y cells, the cells were differentiated in differentiating media containing 10μM of ATRA till 7^th^ day. Following differentiation (referred as day 0), cells were exposed to 50 nM rotenone every alternate day till 14^th^ day and cells were then processed for immunofluorescence and immunoblotting(Borland et al. 2008). All the experiments were repeated at a minimum of three times (represented as ‘n_exp_’ in figure legends) for statistical significance.

### Plasmids, cloning, mutagenesis and transfection

Myc-DDK tagged wild-type human UBA52 (Myc-UBA52), Myc-DDK tagged wild type human α-synuclein (Myc-α-SYN) plasmid constructs were purchased from OriGene technologies, whereas UBA52^K48R^ (Flag-UBA52^K48R^), UBA52^K63R^ (Flag-UBA52^K63R^) mutants were customised and purchased from GenSript. All the plasmids were propagated in DH5α cells, followed by their isolation using commercial plasmid isolation kit (Promega, USA). For ectopic expression of the transients, the expression plasmids were transfected in SH-SY5Y cells using lipofectamine 3000 based on manufacturer instructions. The total RNA was extracted after 24 h of incubation (Verma et al. 2018). The cDNA was obtained by RT-PCR and amplified through PCR using respective primers (OriGene Technologies) followed by agarose gel electrophoresis to assess the gene expression. An empty vector, PcDNA3.1 was used where necessary in order to adjust the DNA amounts.

### siRNA mediated knockdown

Pre-designed siRNA constructs were used for RNA interference, procured from Invitrogen (AM16708; Assay ID 102888) along with a scrambled siRNA (4390843). Transient silencing of UBA52 was done using lipofectamine 3000 as described in the transfection kit (L3000-001).

### Animals and stereotactic neurosurgery

The SNCA C57BL/6J-Tg (Th-SNCA*A30P*A53T) 39Eric/J transgenic mice (Stock number: 008239) were purchased from the Centre of Cellular and Molecular Biology (CCMB), Hyderabad and used in this study as a transgenic mice model [CSIR-CDRI (IAEC/2021/SI no2)] to assess the interaction of UBA52 with α-synuclein in genetically-induced PD (Richfield et al. 2002). The genotyping of 12-month-old mice was done using DNA extracted from their tail tips, followed by the PCR assay [Cycling conditions: 94°C for 5 minutes (94°C for 15 seconds, 65°C for 45 seconds and 72°C for 30 seconds) * 35 and 5 minutes at 72°C obtained from the genotyping protocol database of Jackson Laboratories (Yan et al. 2022). The primers used for genotyping are mentioned in the Suppl. Table-2.

For sporadic neurotoxin-induced experiments, the outbred strain of male rat (Sprague-Dawley) weighing 200-220 g was procured from the National Laboratory Animal Centre of CSIR-Central Drug Research Institute. All conducted experiments followed the strict guidelines of the Institutional animal ethics committee [CSIR-CDRI (IAEC/2018/F-52)]. Four animals were maintained in each polyacrylic cage with food and water ad libitum. Standard ambient conditions with 12h light and dark cycle, room temperature 22±1°C and humidity 60-65% were provided. The rats were divided into two groups having 4 animals per group. The groups were control (sham-operated DMSO-injected) and diseased (rotenone administered sporadic PD). The experimental rats were anesthetized using a mixture of xylazine (10 mg/kg) and ketamine (80 mg/kg), mounted on the stereotaxic apparatus (Stoelting, USA) and a specific region was located by measuring coordinates from bregma(Anon n.d.). Unilateral administration (on the right side of rat brain) of rotenone (dissolved in 3μl DMSO, 6μg in each region) was given in the substantia nigra (SN) and striatum (STR) region (Gupta, et al. 2021). Appropriate care of rats was taken to prevent mortality and after completion of experimental duration (3 days post administration) the rats were sacrificed, both SN and STR were isolated and processed for various assays or the brains were fixed for histological or immunofluorescence studies.

For α-synuclein-PFFs (preformed fibrils)-induced experiments, 4μl of 2μg/μl of recombinant mouse α-synuclein protein PFF (Abcam, ab246002) was unilaterally administered in the striatum region of the rat brain and simultaneously 4μl of PBS was injected in the sham-operated control rats (Paumier et al. 2015). Appropriate care of rats was taken to prevent mortality and after completion of experimental duration (15-, 30- or 45-days post administration of α-synuclein protein PFF) the rats were anesthetized, followed by intracardiac perfusion with 0.9% saline and decapitated to remove the brain quickly (Candelario-Jalil et al. 2000). SN and STR regions were isolated from brain and the tissues were processed for various assays.

All the experiments have been repeated a minimum of three times (represented as ‘n_exp_’ in figure legends) for statistical significance.

### Animal Behaviour Assessment

#### Neuromuscular coordination

Rotarod assay was performed to assess the neuromuscular coordination in the control and rotenone-treated SD rats (Monville et al. 2006). Briefly, the rats were trained for 3 consecutive days on the rotarod apparatus during a habituation trial of 5 minutes at 5-20 rpm to learn the motor balance. Before sacrifice, the rats were subjected to three rotarod sessions at 20 rpm for 2 minutes each. The data was manually recorded for all experimental groups and the frequency to fall was calculated per minute.

#### Stereotype behaviour

Apomorphine (1mg/kg) was given to control and rotenone-administered SD rats by intraperitoneal route to assess the stereotype behaviour. After 30 minutes of apomorphine injection, the rotations were recorded on a live-video system. Rotations were observed for 2 minutes and only full body rotations were counted. Data were articulated as contralateral rotations per minute.

### Cell viability

Cell viability of SH-SY5Y was estimated using mitochondrial dehydrogenase activity assay (tetrazolium dye-based colorimetric test) as reported previously (Gupta et al. 2015). The absorbance was read at 550 nm wavelength by a spectrophotometer (Gen5, BioTek) and mean of optical density was illustrated in graphs.

### mRNA Expression by RT-PCR & qPCR

RT-PCR was performed as reported previously (Verma et al. 2018). For qPCR, the PCR mixture was amplified in a DNA thermal cycler (Applied Biosystems) for 35 cycles and the products were identified on a 2% agarose gel electrophoresis containing ethidium bromide. The respective primers were procured from Integrated DNA Technologies (Suppl. Table-2). Images were captured by an UVI gel documentation system and intensity was measured by Image J software.

### Protein extraction and western blot

The protein was extracted from cell and tissue lysates as reported previously and immunoblotting was performed (Singh et al. 2022) using primary antibodies with appropriate secondary antibodies. The immunoblots were visualized using ChemiDoc XRS+ (Bio-Rad) after developing the signal with substrate Femto Lucent plus HRP (G-Biosciences). Mean intensity (intensity/area) of bands was determined and normalized against ß-actin using ImageJ software (NIH, USA).

### Immunocytochemistry / Immunohistochemistry and counterstaining

#### Immunocytochemistry (ICC)

The SH-SY5Y cells were seeded on the poly-l-lysine coated coverslips and treatment was given for 24 h. ICC was performed in control, treated & transfected cells as reported previously (Verma et al. 2018). The signals were captured through fluorescent (Nikon eclipse E200, Japan) or confocal microscopy (Carl Zeiss).

#### Immunohistochemistry (IHC)

After the treatment rats were sacrificed, brain was perfused with 15-20 ml of PBS. Quickly each brain was isolated and kept in 4% sucrose solution for 3 h, then incubated in 10% sucrose solution for another 3 h, finally shifting the brains to 30% sucrose solution and kept overnight at 4°C. The brains were then stored at −20°C for 2 h, blocks were prepared with cryomatrix (Invitrogen) and sections were cut down using cryostat (Thermo Scientific) of thickness of 10-15 microns. The sections were collected on the poly-l-lysine coated slides immediately and processed for staining. Finally, the sections were mounted using an anti-fade medium containing counter stain DAPI (nuclear stain) and images were captured by fluorescent microscope (Nikon eclipse E200, Japan).

### Thioflavin-S (Th-S) Assay

This assay was performed to assess the protein fibril formation. The cells were seeded on poly-l-lysine coated coverslip, followed by transient expression of Myc-UBA52 and rotenone treatment. Following treatment, the cells were washed in PBS and fixed with 4% PFA in PBS as mentioned above. For staining, the cells were incubated with Th-S stain (0.05% prepared in ethanol) for 30 minutes at room temperature in the dark. The cells were washed thrice with 70% ethanol then once rinse with PBS and mounted on slides using an anti-fade medium with DAPI to visualised under fluorescent microscope (Nikon eclipse E200, Japan).

### Confocal Microscopy

Confocal imaging was performed in SH-SY5Y cells to visualize the localization/co-localization of α-synuclein, UBA52 and HSP90 (Verma et al. 2018). After staining, the cells were mounted in DAPI containing anti-fade mounting medium. The slides were visualized by confocal microscope (Carl Zeiss) and images were captured in a single z-confocal axis.

### Co-immunoprecipitation (Co-IP)

SH-SY5Y cells or dissected rat brain tissue samples were lysed in lysis buffer containing phosphatases and proteases. An equal amount of lysis extracts (1-2 mg) was incubated with anti-UBA52 antibody and protein A-Sepharose beads to make complex with end-over-end rotation overnight at 4°C. Non-specific bound proteins were removed by 4-5 times washing with chilled PBS and the immunocomplexes were boiled with 2x Laemmli buffer at 95°C for 10 minutes to denature the existing proteins. The proteins were resolved on 4-20% gradient SDS-PAGE gel and either immunoblotted with respective antibodies for visualizing the proteins using chemiluminescence or processed to prepare the sample for mass spectrometric analysis.

### Mass spectrometry

Cell or tissue lysates were resolved on a 4-20% gradient pre-cast SDS-PAGE and later incubated in Coomassie blue (G-250) for 2-4 h till the gel was stained followed by de-staining (50: 40: 10:: dH_2_0: Methanol: Glacial acetic acid solution) at room temperature. Once the separated bands became visible, the de-staining solution was removed and individual bands were sliced using a fine blade and immersed in microcentrifuge tubes (MCTs) containing the de-staining solution to remove the stain completely. Next, the gel slices were hydrated in 50-100μl of 25 mM ammonium bicarbonate (ABC) solution (Step 1). This was followed by dehydrating the gel slices in 50 μl of solution A (2:1 mixture of acetonitrile: 50mM ABC) for 5 minutes at room temperature (Step 2). The above steps were repeated till the gel slices became transparent. Next, the gel slices were vortexed for 5 minutes in 100% acetonitrile (ACN) and evaporated on a speed vacuum for 30 minutes at 45°C for complete dehydration. The gel slices were rehydrated with 150 ng of trypsin (Sigma) and incubated on ice for 60 minutes. After the gel slices were completely rehydrated, 50-100μl of 25 mM ABC solution was added and gel slices were incubated overnight at 37°C. Following day, the supernatant was taken in a new MCT and each gel slice was dipped in 20-30 μl of 50% ACN & 2% trifluoroacetic acid (TFA) and vortexed for 5 minutes. All these extractions were added to the MCT containing previous supernatant and evaporated on a speed vacuum till completely dry. The lyophilized sample was resuspended in 30% ACN and 0.1% TFA (5-10μl) and again vortexed for 15 minutes to completely dissolve the sample. Volume of 0.75μl of sample was mixed with equal amount of matrix [α-cyano-4-hydroxycinnamic acid (α-CHCA)] (1:1), spotted on a MALDI plate and processed using AB Sciex 4800 MALDI-TOF/TOF mass spectrometer. Positive ion spectra over m/z 800-4000 Da were recorded for analysis of UBA52 interacting proteins. From each spectrum, maximum of twenty-five precursors with minimum signal: noise ratio was carefully chosen for MS/MS analysis. Representative peptides spectrum against each sample was identified using Protein Pilot (AB Sciex) and corresponding UBA52 interacting proteins were predicted using MASCOT and NCBInr database.

### *In vitro* Ubiquitylation assay

For *in vitro* ubiquitylation of HSP90 and α-synuclein, the cell and tissue lysates were prepared using lysis buffer. Co-immunoprecipitation was performed using anti-UBA52 antibody-protein A-Sepharose beads complex and the immunocomplexes obtained were rinsed with chilled PBS for 4-5 times and loaded on the vial, containing the reaction components provided in *in vitro* ubiquitylation assay kit (Enzo life sciences). The assay was performed based on manufacturer’s instructions at 37°C for 90 minutes and the reaction was terminated after addition of 2x non-reducing Laemmli buffer. The samples were resolved on SDS-PAGE, immunoblotted with anti-HSP90 and anti-α-synuclein antibody followed by their visualization using chemiluminescence.

### Proteasome Activity

Proteasome activity was estimated in both cell (WT and Myc-UBA52) and tissue lysates as reported by us previously (Gupta, Tiwari, et al. 2021) utilizing trypsin (Z-ARR-AMC) and chymotrypsin (SUC-LLVY-AMC) substrate (Enzo life sciences). Reaction mixtures were incubated for 60 minutes at 37°C in dark and the fluorescence intensity was measured using fluorimeter (Varian Cary Eclipse, USA) at excitation/emission 360/460nm, respectively.

### Statistical analysis

Data were analysed using Student’s unpaired t-test or one-way analysis of variance (ANOVA) and the difference between control and treated sets was analysed by post hoc Dunnett’s multiple comparison or Newman Keul’s test. The data generated upon assessing the effect of Myc-UBA52 in neurotoxin-induced studies were analysed using two-way ANOVA, followed by Tukey’s-multiple comparison test. Values are expressed as the mean ± SEM and the p value less than 0.05 was considered as statistically significant.

## Results

### Parkinson’s disease specific pathological markers and UBA52

During the physiological condition, the cellular protein aggregates are degraded through protein degradation mechanisms like UPS which mandatorily requires ubiquitin to process target proteins for their degradation through proteasome machinery. With this hypothesis, we first performed the *in silico* analysis utilizing various database platforms like BGEE (*https://bgee.org/*) and NCBI-Geoprofile (*https://www.ncbi.nlm.nih.gov/geoprofiles/*) to check the available data of ubiquitin encoding genes in PD related brain regions. BGEE database suggested high basal expression of UBB, UBC, UBA52 and RSP27a genes in PD-related brain region-substantia nigra (SN). Next, we explored various publicly accessible microarray data available in the geoprofiles to observe the expression level of ubiquitin genes [UBB, UBC, UBA52, RPS27a] in human brain (control and PD) as well as in rat brain (depending on available database). We identified a geoprofile, GDS2821 showing the low abundance of UBA52 in PD *post mortem* brain in comparison to healthy control human *post mortem* brain. In addition, we also identified the geoprofile, GDS5646 showing the altered level of UBB, UBC, UBA52 and RPS27a in peripheral blood. A microarray-based gene expression profiling done in post-mortem substantia nigra of the human brain of control and PD subjects showed the significant depletion of UBB (5.9 folds) and UBA52 (2.1 folds) in the PD brain (Simunovic et al. 2009) However, the role of UBB in neurodegenerative disease is well studied (Gentier & van Leeuwen 2015; Lindsten et al. 2002; Fischer et al. 2003) we, therefore performed studies to check alteration in gene expression and protein level of UBA52 during to the onset of PD.

Based on the *in silico* inference and previous preliminary findings, we conducted the experiments to validate the alteration of UBA52 in PD-specific pathological markers in both sporadic cellular and rat experimental models and SNCA C57BL/6J-Tg (Th-SNCA*A30P*A53T) 39Eric/J transgenic mice.

Rotenone is a potent toxin that cause mitochondrial complex I inhibition, selectively leading to nigrostriatal dopaminergic neurodegeneration and motor deficit (Sherer et al. 2003; Alam & Schmidt 2002; Betarbet et al. 2000). In contrast to 6-OHDA and MPTP, it is also shown that the rotenone injected SNpc region contains proteinaceous inclusions like the Lewy bodies, immunoreactive for ubiquitin and alpha-synuclein (Sherer et al. 2003).

In line with this evidence, the selection was done and first, the dose-ranging was performed to observe the appropriate concentration of rotenone that causes PD-related diseased condition along with cell viability assessment in SH-SY5Y cells (Fig 1a-b). A significant decrease in cell viability along with depletion in mRNA level of tyrosine hydroxylase (TH) was observed at both 250nM and 500nM doses of rotenone. Next, we evaluated the stability of PD pathology in the employed rotenone-induced experimental rats by estimating the mRNA levels of TH at different time points (till 21 days of rotenone administration) in both the SN and STR regions of brain. Data suggested persistent depletion of TH in experimental rat model till 21 days which was initiated at 3^rd^ day after rotenone administration (Fig 1c). Findings are in concordance with the study of Faull & Laverty (1969) and Bové et al. (2005) that showed the maximal depletion of dopaminergic neurons at 3 days post neurotoxin administration. This might be due to the employed dose as well as the sites of injection (both SN and STR) of rotenone in the rat brain. We therefore, selected the 3-day time point to study intracellular signaling mechanisms at early/initiatory phase of disease. The behavioural parameters (apomorphine-induced rotations and rota rod assay) also showed significant impairment in neuromuscular coordination after 3-day of disease induction (Fig 1d-e). Concurrently, we utilised the same cDNA samples [used to assess the TH mRNA] to check the expression of ubiquitin genes. Experiments confirmed the depletion in mRNA level of UBA52 in both substantia nigra (SN) and striatum (STR) regions of rat brain (Fig 1f), whereas the level of UBB, UBC and RPS27a was apparently not altered (Suppl. Figure-1). Simultaneously, we observed the low abundance of UBA52 mRNA in neuronal cell line SH-SY5Y after treatment (Fig 1g). In line with above findings, we further estimated the protein level of UBA52, TH, α-synuclein and cleaved caspase-3 in both SH-SY5Y cells and in SN & STR regions of rat brain. Data showed significant alteration in protein level of UBA52, TH, α-synuclein and cleaved caspase-3 after rotenone administration in both experimental models (fig 1h-m). However, in SH-SY5Y cells the alteration in protein level of TH and UBA52 was significantly profound at 500nM concentration (Fig 1f) therefore, further experiments were conducted at 500nM concentration of rotenone.

**Figure 1:**
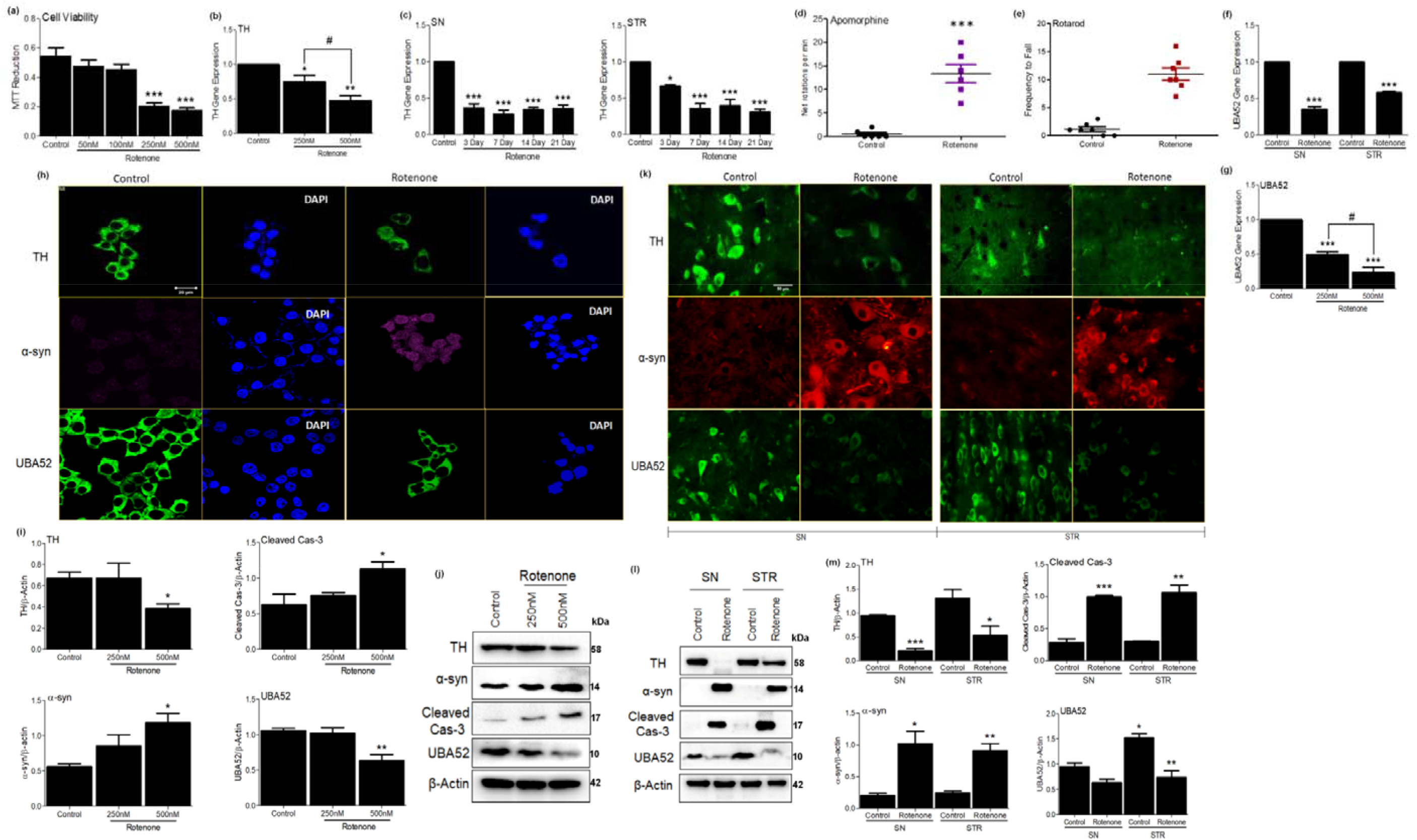
Parkinson’s disease markers and UBA52 expression in SH8Y5Y and rat brain. **(a)** Graph representing MTT reduction assay in SH-SY5Y cells to confirm optimum treatment dose of rotenone. **(b)** q-PCR of TH in SH-SY5Y cells and **(c)** SN (substantia nigra) & STR (striata) region of the rat brain, normalised against β-Actin to determine the time-point of study and rotenone concentration in *in vitro* and *in vivo* model; n_exp_=3. **(d & e)** Behavioural study of control and rotenone-lesiored rat. Contralateral rotations induced by apomorphine **(d)** and lapse in muscle-coordination measured through **(e)** rotarod test ; n_exp_=3. **(f)** q-PCR of UBA52 in control and nearotoxin-treated rat brain regions, normalised against β-Actin; n_exp_=4. **(g)** q-PCR of UBA52, normalised against β-actin in SH-SY5Y cells after rotenone treatment n_exp_ =3. **(h)** Confocal microscopy images depicting expression of TH, α-syn and UBA52 after secondary labelling with Alexa fluor - 488 (green) and -546 (red). −647 (pink) in control vs rotenone treated SH-SY5Y cells, Scale bar; 20μm; n_exp_=3. **(i & j)** Graphical and pictorial illustration of western blot performed to assess the expression of TH, α-syn cleaved cas-3 and UBA52, normalised against β-Actin in human SH-SY5Y neuroblastoma cells; n_exp_·3. **(k)** Flourescent images depicting the expression of TH, α-syn and UBA52 after secondary labelling with Alexa fluor −488 (green) and −546 (red) in control vs rotenone treated SH-SY5Y cells Scale bar; 50μm; n_exp_=3. **(I)** Immunoblot images illustrate the protein level of TH, α-syn, cleaved cas-3 and UBA52, normalised against β-Actin in rat brain regions. **(m)** Graphical representations of western blot images normalised against β-Actin in SN and STR rat brain regions indicating the similar pattern as observed in SH-SY5Y cells; n_exp_=3. Quantiflication are mean and SEM of at least three independent experiments and statistical analysis were performed using Student’s t-test and one-way ANOVA, followed by Dunnett’s post hoc test. * p<0.05, ** p<001, *** p<0.001 control vs. Rotenone; # p<0.05 R250nM vs. R500nM.

### UBA52 surplus attenuates the PD specific pathological markers and neuronal death

Having determined that UBA52 was decreased in both *in vitro* and *in vivo* rotenone-treated experimental models, we then sought to validate the role of UBA52 in alteration of disease specific pathological markers. Primarily, *in silico* analysis indicated the interaction of UBA52 with TH and α-synuclein as assessed through a freely accessible server, PSOPIA (Prediction Server of Protein-Protein Interaction) based on both human and rat protein sequences available on UniProtKB database (Murakami & Mizuguchi 2014). Results indicated that UBA52 interacts with α-synuclein (Sall-0.9123) to a greater extent in comparison to TH (Sall-0.3613) (Suppl. File-1). Further, to confirm the interaction of UBA52 with α-synuclein, we performed co-immunoprecipitation in both utilized rotenone-treated models. Observations validated *in silico* finding and showed that PD induction enhanced the interaction of UBA52 and α-synuclein. It might be because UBA52 is a ubiquitin gene and it interacts with misfolded proteins or proteins with increased abundance, during disease pathogenesis to enhance their proteasome-mediated degradation and restore homeostasis (Fig 2a-b).

**Figure 2:**
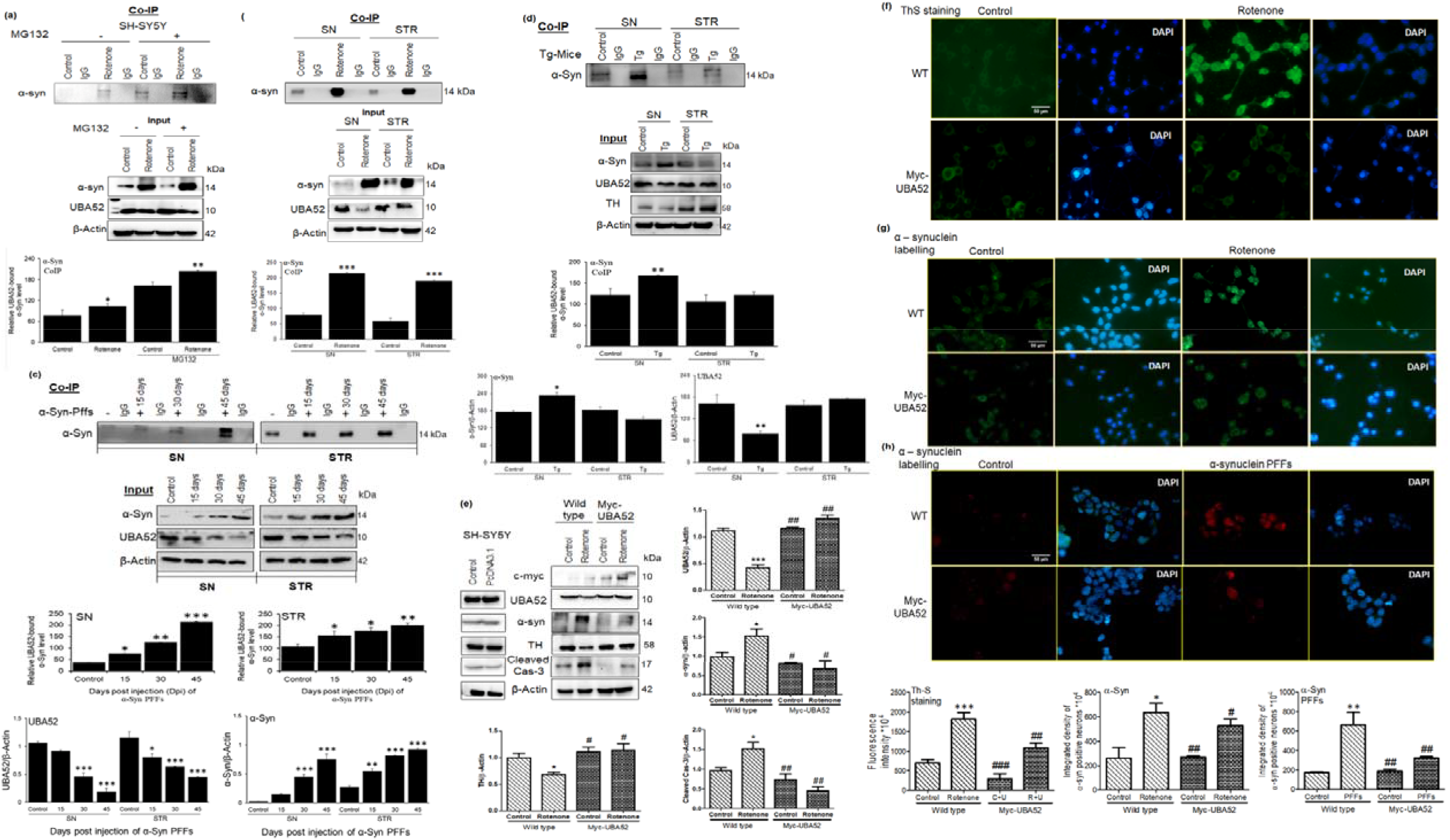
UBA52 interacts with α-Syn and its over-expression in SH-SY5Y cells deter the alteration in PD markers protein abundance, preventing the disease onset. **(a,b)** Representative images and analysis depicting the CoIP experiment to check *in vitro* interaction of UBA52 and α-Syn against IgG as a negative control in SH-SY5Y cells with or without rotenone& MG132 treatment, and control& rotenone treated PD-related SN and STR region, respectively, rats used per group in each replicate=4. n_exp_=4. **(c)** Representative image and analysis depicting the CoIP experiment to check the interaction of UBA57 and α-Syn, against IgG as a negative control in PRS-admintstered control& mouse recombinant α-Syn PFF treated PD-relaled SN and STR region, respectively at 15-, 30- and 45-days poet injection (dpi) to show the underlying mechanism of UBA52 in regulating α-Syn abundance; rats used per group per replicate=5. n_exp_=3. **(d)** Representative image and analysis depicting the CoIP experiment to check the interaction of UBA52 and α-Syn, against IgG as a negative control in the PD-related SN and STR region of the control C57BL mice and the C57BL/6J-Tg (Th-SNCA-A30P*A53T) 39Eric/J transgenic (Tg) mice, respectively mice used per group per replicate=5. n_exp_=2 **(e)** PcDNA3.1 or Myc-UBA52 transfected SH-SY5Y cells with or without rotenone treatment were immunoblotted for PD-related protein markers; n_exp_=3. **(f)** Thioflavin-S (Th-S) staining and the fluorescence intensity of Th-S positive neurons to detect protein aggregates in wild-type or Myc-UBA52 transfected SH-SY5Y cells with or without rotenone treatment. Scale bar-50μm; n_exp_=3. **(g)** Florescent images and the integrated density depicting cytological staining for α-Syn positive neurons, later tagged with Alexa fluor-488 (green) in wild-type or Myc-UBA52 transfected SH-SY5Y cells with or without rotenone treatment Scale bar-50μm; n_exp_=3. **(h)** Florescent images and the integrated density depicting cytological staining for α-Syn positive neurons, later tagged with Alexa fluor-532 (red) in wild type or Myc-UBA52 transfected SH-SY5Y cells with or without human recombinant α-Syn PFF treatment. Scale bar-50μm; n_exp_=3. Quantification is the mean and sem of at least three independent experiments and statistical analysis was performed using two way-ANOVA, followed by Tukey’s multiple comparison test * p<0.05. ** p<0.01, *** p<0.001 control vs. Rotenone/αSyn PFFs/Transgenic (Tg); # p<0.05, ## p<0.001, ## p<0.001 Myc-UBA52 vs. Rotenone.

α-synuclein pre-formed fibrils (PFFs) efficiently participate to seed the aggregation and fibril formation of the endogenous α-synuclein. Therefore, to check the *in vivo* relevance or underlying mechanism for the direct UBA52-dependent regulation of α-synuclein via interaction, we performed the co-immunoprecipitation study in the SN and STR regions of recombinant mouse α-synuclein protein PFF-injected SD rats. Consistent with the hypothesis, we observed a time-dependent increase in the interaction of UBA52 and α-synuclein in the striatum (Fig 2c) along with the time-dependent altered protein levels of both UBA52 and α-synuclein (Input bands). However, in the SN region, the interaction was observed 30 days post injection (dpi) which profoundly increased at 45dpi (Fig 2c). The protein abundance of UBA52 was visibly reduced from 30dpi to 45dpi opposite to the change in α-synuclein protein which increased from 30dpi to 45dpi (Input bands).

Next, the interaction of UBA52 and α-synuclein was further assessed in the Tg-SNCA mice to confirm their interaction in the mutant α-synuclein model. Data obtained showed that this interaction was visibly higher in the SN region of the Tg-SNCA mice in comparison to the control C57BL mice (Fig 2d). The protein level of UBA52 was reduced in the Tg mice, whereas, α-synuclein protein level increased in the SN region of the mice. However, in the STR region, we did not observe a stark difference in the interaction of UBA52 and α-synuclein as well as in the protein abundance of UBA52 and α-synuclein (Fig 2d). It may be due to the middle-age (12 months) of the transgenic mice, wherein the observed changes occurred in the SN region and the old-aged mice (15-23 months) would be useful for further establishing the interaction profile of UBA52 and α-synuclein in the STR region of the brain. Parallelly, we also found only slight depletion in TH protein level in the SN region with a rather increased TH abundance in the STR region, confirming the previous reports (Chesselet 2008; Dawson et al. 2010) which showed that α-synuclein mutation using transgenic approaches doesn’t elicit the neurodegeneration in DA neurons.

Further, the SH-SY5Y cells were transiently overexpressed for UBA52 and we observed that UBA52 overexpressed cells exhibited significant attenuation of rotenone-induced upregulated α-synuclein and TH depletion along with inhibited neuronal death (cleaved caspase-3 level) (Fig 2e). Altered protein folding and degradation during stress conditions is one of the most studied phenomena in PD pathology (Tiwari & Singh 2020) therefore, next the protein fibril formation corresponding to aggregate generation was assessed through thioflavin staining. In agreement with this, UBA52 overexpressed cells showed inhibited fibril formation in comparison to wild-type cells during the early time-point of disease induction (Fig 2f). Since α-synuclein is the major protein associated with PD and a prime component of the Lewy body along with chaperones and ubiquitin, we further assessed the level of α-synuclein by immunostaining. Rotenone treatment in the SH-SY5Y cells significantly exhibited the augmented level of α-synuclein which was suppressed after Myc-UBA52 overexpression in SH-SY5Y cells (Fig 2g). The implication of UBA52 transient overexpression was also confirmed in the recombinant human α-synuclein protein PFF-treated SH-SY5Y cells, and the immunofluorescent images suggested that UBA52 overexpression prevented the tendency of human α-synuclein protein to form the toxic fibrils (Fig 2h). These data revealed the *in vitro* and *in vivo* relevance for the direct UBA52-dependent regulation of α-synuclein via interaction along with the strong implication of UBA52 in regulating the PD-specific pathological markers and protein aggregation tendency during the acute phase of disease pathogenesis.

### UBA52 revokes the PD specific pathophysiological markers in differentiated neuronal cells

Upon obtaining significant results in bridling the effect of transient overexpression of Myc-UBA52 for PD-specific pathological markers (TH and α-synuclein) in an experimental acute model of SH-SY5Y, we next studied the effect of UBA52 in reversing the rotenone-induced PD conditions in differentiated neuronal cells employing SH-SY5Y, N2a and PC12 cells. Cells when cultured in a respective differentiating medium, showed high expression of differentiating markers (Tuj1 and Map2) and dopaminergic neuronal marker (TH) in SH-SY5Y, N2a and PC12 cells (Suppl. Figure-2, Fig 3a). To note, ATRA treatment did induce some cell death in all the neuronal cells as reported previously in various differentiation-inducing protocol studies in contrast to NGF treatment in PC12 cells. After 7 days of differentiation in respective cell lines, we investigated the effect of UBA52 on PD-specific marker TH, α-synuclein and neuronal apoptosis (cleaved caspase-3). Analysis of immunoblots in both wild-type and UBA52 overexpressed neuronal cells revealed the significant preventive effect of UBA52 in rotenone-induced diseased conditions as assessed by protein level of TH, α-synuclein and cleaved caspase-3 (Fig 3b-d). Although ATRA has not been reported as the differentiation-inducing chemical in N2a cells to promote dopaminergic characteristics, astonishingly, our results showed an apparently high expression of TH after 7-days of ATRA treatment (Fig 3d).

**Figure 3:**
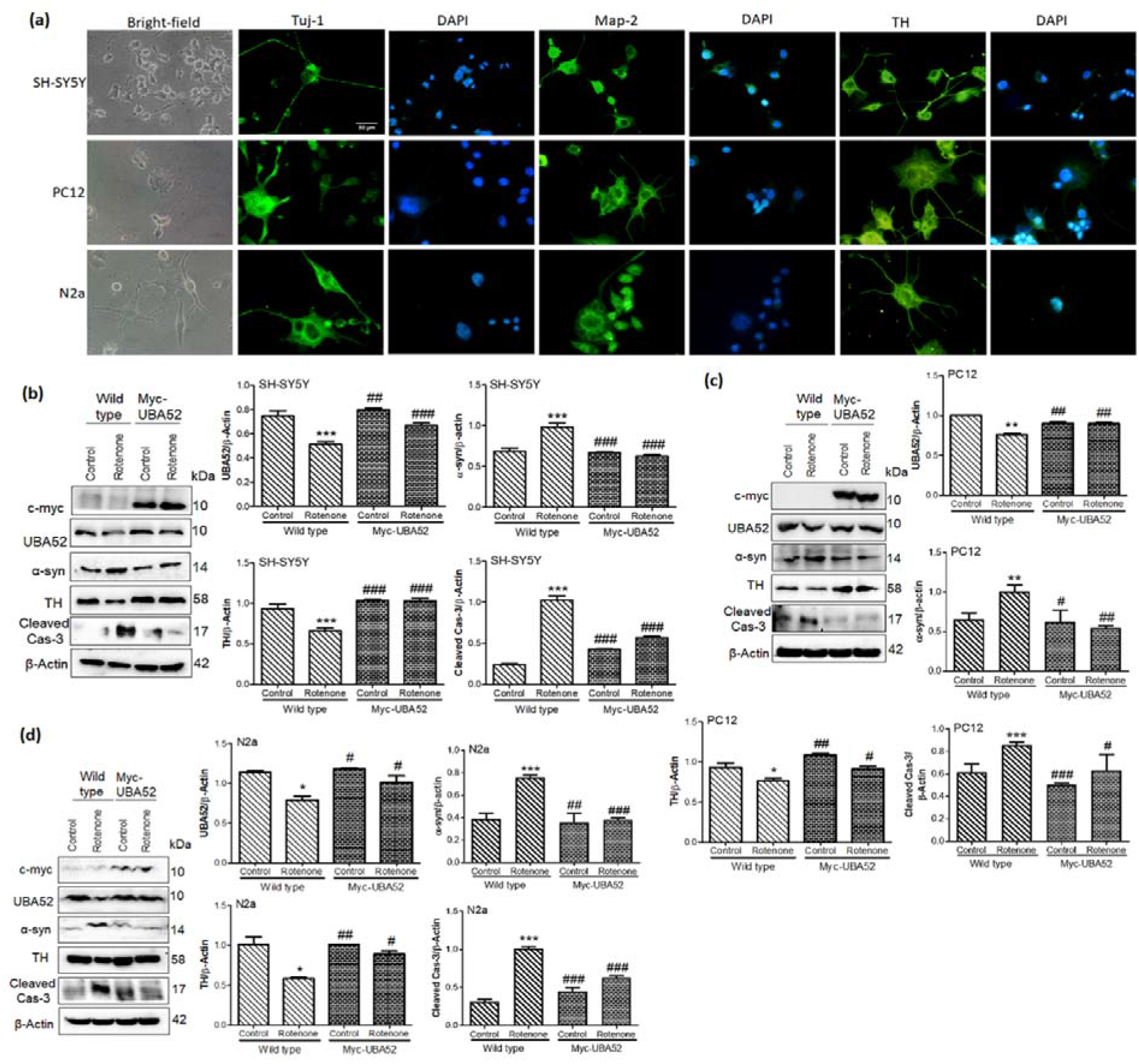
Effect of UBA52 overexpression on PD-related pathological markers In differentiated neuronal cell lines. **(a)** Representative image depicting the bright-field (20x) and immunofluorescence (40x) of differentiation markers Tuj-1, Map-2 and dopaminergic marker tyrosine hydroxylase (TH) in SH-SY5Y. PC12 and N2a cells, Scale bar- 50μm. **(b, c and d)** Blots and graphical representations illustrating the PcDNA3.1 or Myc-UBA52 transfected SH-SY5Y PC12 and N2a differentiated cells to show the PD-related protein markers expression with or without rotenone treatment n_exp_=3. Quantification are mean and SEM of at least three independent experiments and statistical analysis were performed using two way ANOVA, followed by Tukey’s-multiple comparison test * p<0.05 ** p<0.01 *** p<0.001 control vs. Rotenone; # p<0.05, ## p<0.01, ### p<0.001 Myc-UBA52 vs. Rotenone.

### Effect of UBA52 on PD-specific pathological markers in experimental *in vitro* chronic model

The study was further expanded to assess the effect of UBA52 on chronic experimental model of PD employing SH-SY5Y cells (Borland et al. 2008). To aim this, we differentiated the SH-SY5Y cells for 7 days and then exposed the differentiated dopaminergic cells to rotenone (50nM) for 14 days (12days rotenone+48h of Myc-UBA52 transient expression) to develop chronic PD *in vitro* model. Rotenone treatment induced the neurite swellings from day-5 with gradual decrease in neurite processes and cell viability till day-14 (data not shown). Immunoblotting data suggested significant alteration in protein level of TH, α-synuclein and cleaved caspase-3 level after rotenone treatment in chronic *in vitro* SH-SY5Y cells, which was discernibly reduced upon transient expression of Myc-UBA52 with or without rotenone exposure (Fig 4a). Cleaved caspase-3 protein level was also observed in control and Myc-UBA52 expressing cells which was accounted for both long duration maintenance of SH-SY5Y cells and initial 7 days of ATRA treatment however, it was less in comparison to treated cells (Fig 4a). In parallel, we also confirmed our findings through immunofluorescence of TH and α-synuclein in chronic *in vitro* model. Similar to immunoblotting, chronic rotenone treatment reduced the neurite processes in comparison to control cells which was relatively less in cells transiently expressing Myc-UBA52 with or without rotenone exposure (Fig 4b). Due to severe loss of cell viability, as evident through observed cleaved caspase-3 level in control cells, the study was restricted to 14 days only. The observed findings in chronic PD-conditions are in concurrence to findings of acute model and suggests that UBA52 is a relevant pathophysiological regulator during PD.

**Figure 4:**
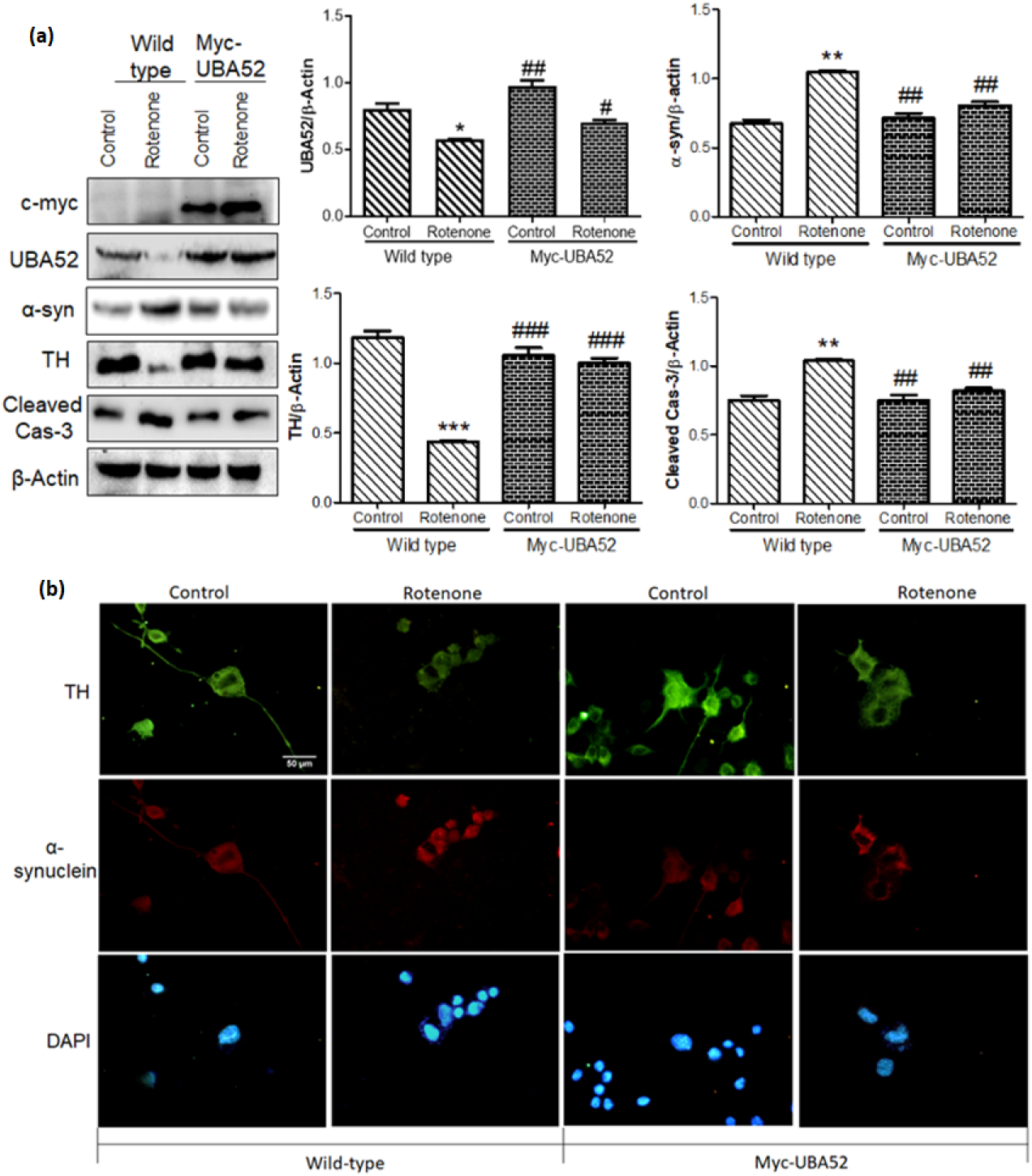
Effect of UBA52 overexpression on PD related pathological markers in chronic *in-vitro* model of SH-SY5Y cells. **(a)** Blots and graphical representations illustrating the PcDNA3.1 (control) or Myc-UBA52 transfected SH-SY5Y differentiated cells with or without chronic exposure of rotenone(50μM) to show PD-related protein markers expression; n_exp_=3. **(b)** Representative image depicting immunofluorescence(40x) of TH and α-synuclein in PcDNA3.1 (control) or Myc-UBA52 transfected SH-SY5Y differentiated cells with or without chronic exposure of rotenone(50μM) to show alteration in neuronal morphology, neurite length and expression of PD-related proteins, Scale bar- 50μm; n_exp_=2. Quantification are mean and SEM of at least three independent experiments and statistical analysis were performed using two-way ANOVA, followed by Tukey’s multiple comparison test. ** p<0.01, *** p<0.001 control vs. Rotenone; # p<0.05, ## p<0.01, ### p<0.001 Myc-UBA52 vs. Rotenone.

### Tuning the interaction between HSP90 and CHIP by UBA52

In view of transiently overexpressed UBA52 mediated neuroprotection in diseased conditions, we searched for the novel interacting partners of UBA52 that might act as a substrate for ubiquitylation in the presence of putative E3 ligases. To accomplish this, we performed MALDI-TOF (matrix assisted laser desorption ionization- time of flight) based mass spectrometric (MS) analysis, with an aim to detect the UBA52-interacting proteins in both cell and tissue lysates. Interestingly, in the series of hits obtained and data analysed by MASCOT software suggested the interaction of various proteins with UBA52. However, this study was focused on the interacting chaperones owing to their critical role in protein degradation mechanisms and henceforth, the data were sorted accordingly. Findings suggested strong interaction of UBA52 with HSP90 (Fig 5a-c; Suppl. Figure:3), which was validated additionally by co-immunoprecipitation using anti-UBA52 antibody and IgG as control. Previously, it has been reported that HSP90 is ubiquitylated by E3 ubiquitin ligase CHIP (carboxy terminus of HSC70 interacting protein) and degraded by proteasome (Kundrat & Regan 2010) therefore, we checked the association of UBA52 and CHIP. Co-immunoprecipitation of lysates using anti-UBA52 antibody showed the high interaction of CHIP after rotenone treatment, suggesting considerable participation of UBA52 in HSP90 ubiquitylation during disease induction (Fig 5d-f).

**FIGURE 5:**
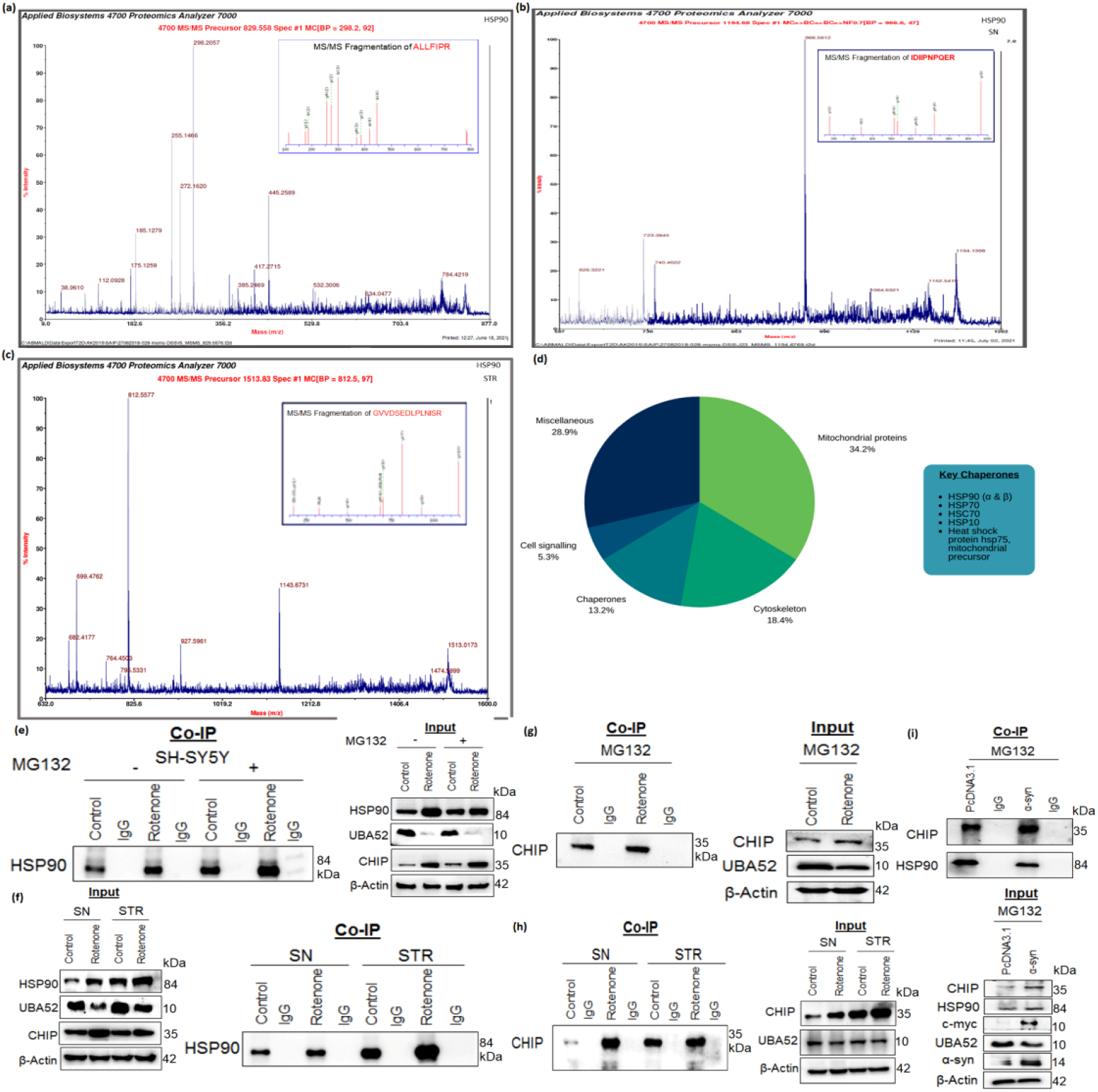
UBA52 interacts with HSP90 and CHIP. MS/MS peaks representing the fragmentation spectrum (highest intensity) obtained from the tryptic peptides of amino acids (331 −337) of HSP90 α/β in **(a)** SH-SY5Y cells and **(b & c)** in rat brain regions; n_exp_=3. **(d)** Pie diagram illustrate the key chaperones obtained from the mass-spectrometric analysis, **(e)** Representative images illustrating co-immunoprecipitation (Co-IP) of UBA52 and HSP90, against IgG as negative control with or without rotenone & MG132 treatment in SH-SY5Y cells; n_exp_=3 and in control and **(f)** rotenone-lesioned SN and STR rat brain region; n_exp_=4. **(g)** SH-SY5Ycells with or without rotenone treatment with MG132. representing the interaction of UBA52 and CHIP along with input bands n_exp_=3. **(h)** Blots representing the interaction of UBA52 and CHIP in control and rotenone treated rat brain regions, substantia nigra (SN) and striata (STR) n_exp_=3. **(I)** Further, the SY-SY5Y cells were transiently overexpressed with Myc-α-synuclein (α-syn) to mimic the diseased condition and the interaction of UBA52 with CHIP & UBA52 with HSP90 was assessed against IgG as negative control in PcDNA3.1 or Myc-tagged α-syn transfected SH-SY5Y cells after MG132 treatment; n_exp_=3.

Further to confirm the specificity of UBA52 in diseased conditions, the SH-SY5Y cells were transiently overexpressed with wild-type α-synuclein (Myc-α-SYN) to imitate the diseased conditions and interaction of UBA52 with HSP90 and CHIP was evaluated. Overexpression of α-synuclein in neuronal cells significantly enhanced the interaction of UBA52 with HSP90 and CHIP (Fig 5g). Simultaneously, we also observed the decreased level of UBA52 (input band) after α-synuclein overexpression (Fig 5g), reaffirming our data of depleted UBA52 in diseased conditions.

Last two years findings also highlighted that both α-synuclein and HSP90 are relocalized to the mitochondrion within the neuronal cell during PD progression (Burmann et al. 2020). Since UBA52 is ubiquitously expressed in the cytoplasm of the neuronal cell, we checked whether it co-localizes with HSP90 and α-synuclein along with its own translocation to other cellular organelles / membranes within the cell following disease induction. Confocal microscopy imaging in SH-SY5Y cells suggested that disease onset enables UBA52 localization in the mitochondrion (Fig 6a-b). Findings also revealed that UBA52 was highly co-localized with HSP90 and α-synuclein in both rotenone-treated and α-Syn PFF-induced SH-SY5Y cells in the cytosol as well as the mitochondrion in comparison to the control cells (Fig 6a-b). Last, we also validated our above findings of SH-SY5Y cells and checked the colocalization of UBA52 with HSP90 and α-synuclein in both SN and STR regions of the rat brain. Microscopic examination revealed a very strong co-localization of UBA52 with α-synuclein as well as with HSP90 in rotenone-induced PD model of rat in comparison to respective control (Fig 6c-d). Altogether, these approaches confirmed high interaction of UBA52 with the molecular chaperone HSP90, E3 ligase CHIP and unequivocally linked PD marker α-synuclein.

**Figure 6:**
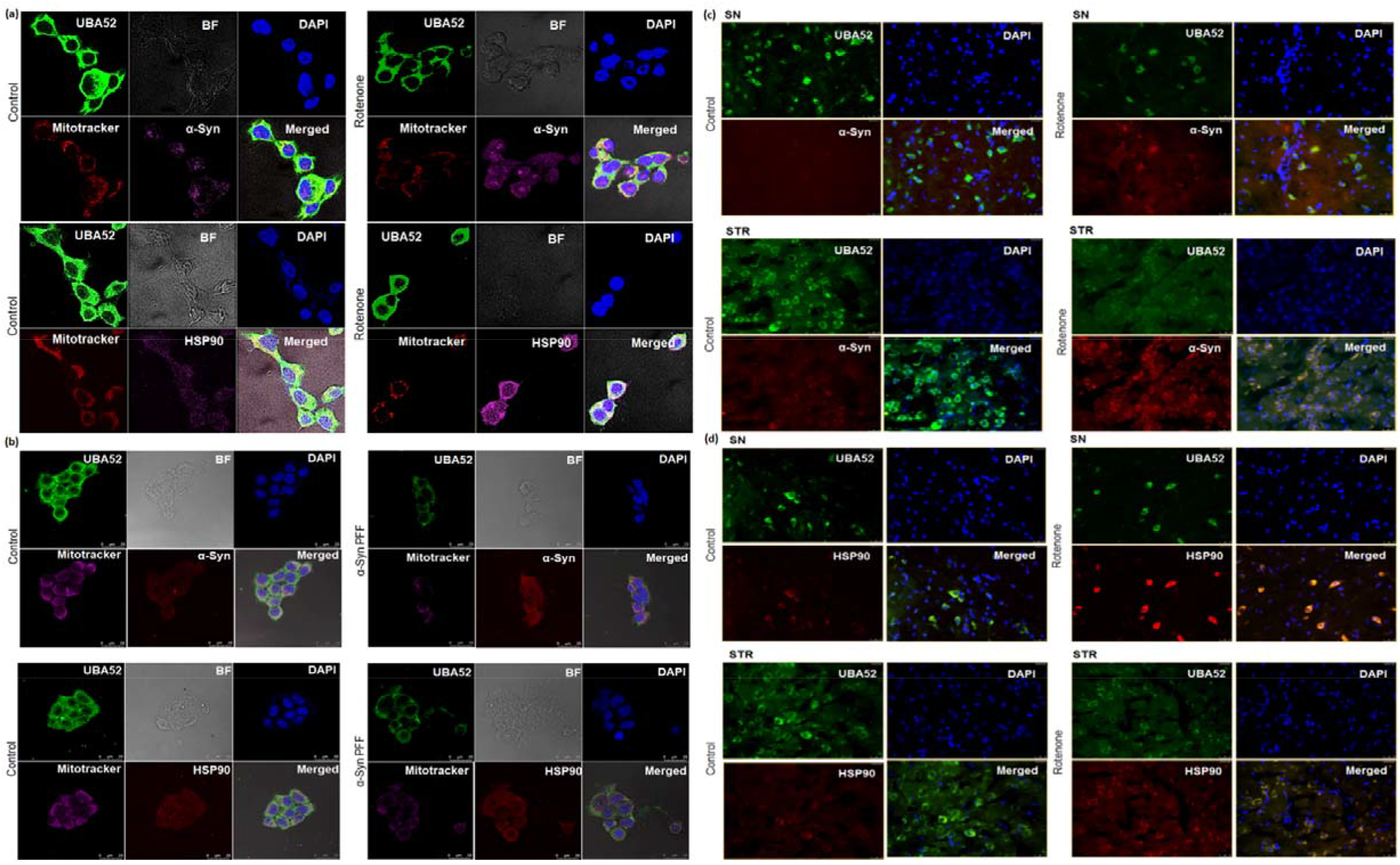
UBA52 colocalize with α-syn and HSP90. **(a)** Confocal microscopy images represent the colocalization of UBA52 and α-syn/HSP90 in control and rotenone-treated SH-SY5Y cells along with the mitotracker deep red (647nm) and DAPI (405nm) representing mitochondrion and nucleus respectively. Alexa fluor: Green-488. Pink-546 ; n_exp_=3 **(b)** Confocal microscopy images represent the colocalization of UBA52 (Alexa fluor Green-488) and α-syn/HSP90 (Alexa fluor Red-546) in control and human recombinant α-synuclein preformed fibrils (PFFs)- induced SH-SY5Y cells along with the mitotracker deep red (647nm) and DAPI (405nm) representing mitochondrion and nucleus respectively. Scale bar- 25μm; n_exp_=2. **(c)** Fluorescent images of rat brain sections representing the colocalization of UBA52 (Alexa fluor Green-488) and α-Syn (Alexa fluor Red-546) in both SN and STR regions, n_exp_=3. **(d)** Fluorescent images of rat brain sections representing the colocalization of UBA52 (Alexa fluor Green-488) and HSP90 (Alexa fluor Red-546) in both SN and STR regions; n_exp_=3.

### Role of UBA52 in HSP90 ubiquitylation during Parkinson’s disease

Since high interaction of UBA52 with HSP90 and CHIP was observed in diseased conditions, we next investigated the role of UBA52 in HSP90 ubiquitylation in consideration of the observed increased level of HSP90 during diseased condition and probable therapeutic implication of HSP90 in PD (Alam et al. 2017). *In vitro* ubiquitylation assay in cells and in rat brain lysates suggested the increase in total ubiquitylation of HSP90 after rotenone administration along with increase in the protein level of CHIP (Fig 7a-b). Also immunoblot probed with anti-HSP90 showed that in SN, the ubiquitylation was apparently more in comparison to the striatum region (Fig 7b) and finding is in concordance to the location of cell body in SN and nerve terminals in the STR. *In vitro* ubiquitylation assay was performed using various E2 enzymes such as UBCH1, UBCH3, UBCH5c, UBCH7, UBCH10 and UBCH13. The E2 enzymes, UBCH5c and UBCH13 tag the protein at position K48 and K63 respectively (Dong et al. 2011). Of all the used E2 enzymes, UBCH13 showed its relevant involvement in ubiquitylation. Since, ubiquitin labelling to target protein takes place through lysine residue, further investigation was performed to assess the specific lysine residue involved in ubiquitylation of HSP90. To attain this aim, we transfected the SH-SY5Y cells transiently with vector encoding Flag-UBA52^K48R^ or Flag-UBA52^K63R^ mutants separately. Immunoprecipitation followed by ubiquitylation analysis showed that K63 mutation in UBA52 prevented the attachment of ubiquitin linkage on the HSP90 protein which led to no change in total ubiquitin as well as total ubiquitylation of HSP90 in comparison to the results generated after mutation in K48 residue of UBA52 (Fig 7c). However, we did not spot similar results of ubiquitylation with α-synuclein as the target protein, confirming the findings from Tofaris et al. (2011) that CHIP and UBA52 (our findings) do not attach ubiquitin chain on α-synuclein in the presence of either UBCH5 (K48 linked) or UBCH13 (K63 linked) E2 ligases. Taken together, our data strongly suggested that E3 ubiquitin-protein ligase CHIP ubiquitylates the target substrate HSP90 through E2 enzyme UBCH13, attaching K63 linked chains in the presence of ubiquitin gene UBA52 (Fig 7d).

**Figure 7:**
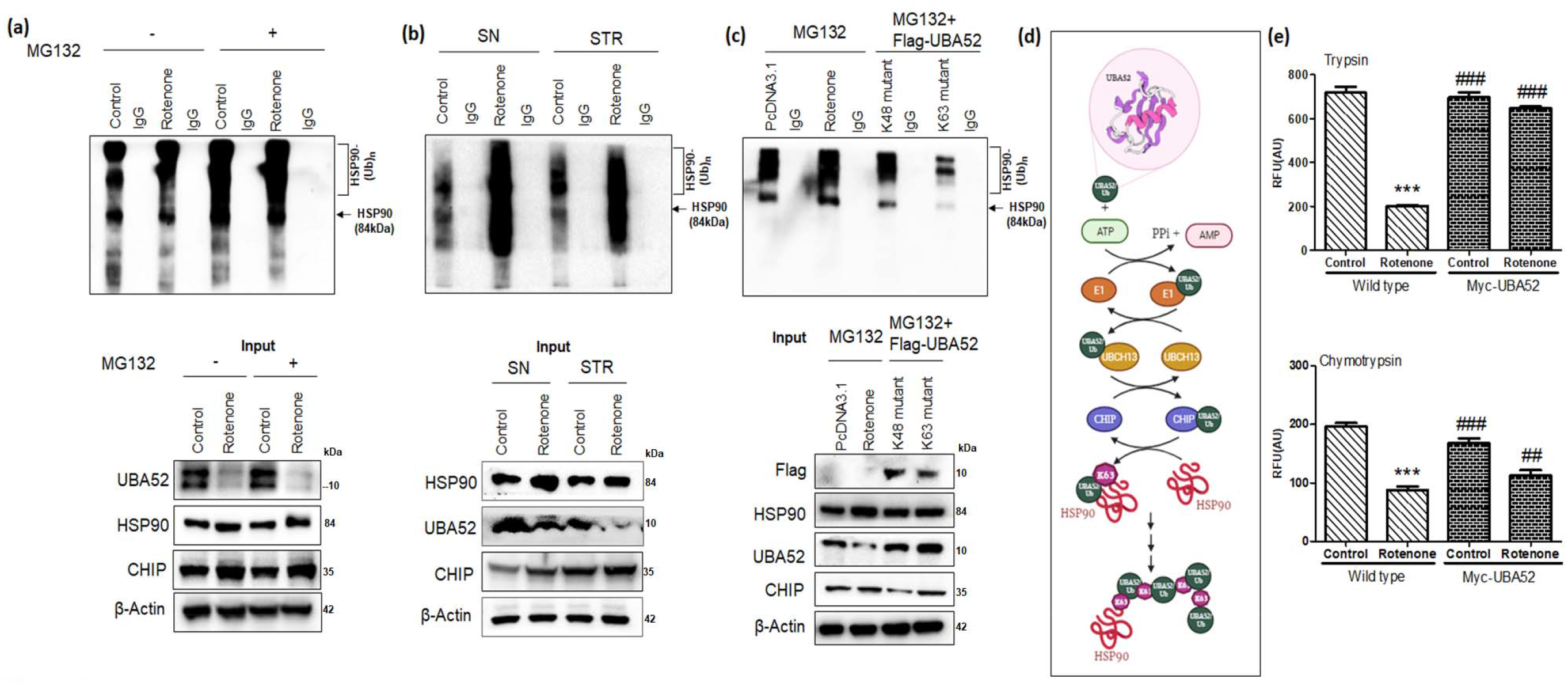
UBA52 is crucial for CHIP mediated poly-ubiquitylation of HSP90 at lysine 63. **(A)** Representative image of *in vitro* ubiquitylation assay after Co-IP of cell lysate of SH-SY5Y cells with UBA52, against IgG negative control with or without MG132 & rotenone treatment; n_exp_=2 **(B)** Immunoblots represent the *in-vitro* ubiquitylation assay after Co-IP of brain lysate of control & rotenone-lesioned SN and STR region of rat brain; n_exp_=2. The positive ubiquitylation reaction was observed in the presence of E2 enzyme, UBCH13 in both the experimental studies. Final reaction mixture was resolved on 4-16% gradient SDS-PAGE and immunoblotted using anti-HSP90 antibody. (C) Next, the SH-SY5Y cells were transiently expressed with mutant UBA52 (Flag-UBA52^K4BA^ or Flag-UBA52^K63A^) to elucidate the specific lysine involved in ubiquitylation of HSP90 after MG132 treatment in control / rotenone treated cells, against IgG negative control; n_exp_=2. (D) Pictorial representation of CHIP-UBA52-UBCH13 mediated K63-linked ubiquitylation of HSP90. (E) Graphical representation for proteasome (trypsin and chymotrypsin) activity in wild type SH-SY5Y cells and after transient overexpression of UBA52 (Myc-UBA52) with or without rotenone treatment; n_exp_=3. Quantification are mean and SEM of at least three independent experiments and statistical analysis were performed using two way-ANOVA, followed by Tukey’s multiple comparison test. *** p<0.001 control vs. Rotenone; ## p<0.01, ### p<0.001 Myc-UBA52 vs. Rotenone.

Protein aggregates accumulate mainly due to decrease in cellular degradation efficiency and decreased proteasome activity with ageing or disease onset (Tai & Schuman 2008). We therefore, checked the proteasome activity in neuronal cells and in rat brain regions during PD condition. Data indicated significant suppression of proteasome activity during diseased conditions in both SH-SY5Y cells (Fig 7e) as well as in SN and STR regions of rat brain (Suppl. Figure-4). Subsequently, we transfected the neuronal cells with Myc-UBA52 and estimated its effect on the proteasomal (trypsin and chymotrypsin) activity. Findings suggested that wild-type UBA52 overexpressed cells resisted the decrease in proteasome activity upon neurotoxin exposure and this observation was starker in trypsin as compared to chymotrypsin activity (Fig 7e). In another set of experiment, we checked the alteration in total ubiquitin levels through immunoblotting after UBA52 knockdown in control SH-SY5Y cells as well as in rotenone-treated cells. Rotenone treatment in neuronal cells increased the total ubiquitin level, whereas, transient transfection of siRNA-UBA52 in control cells led to depletion in total ubiquitin levels. The upregulated ubiquitin level in rotenone-treated cells might be due to the activation of various downstream neurodegenerative mechanisms which together increases the cargo load on the UPS to clear the aggregate-prone proteins, thereby raising the total ubiquitin level during early phase of disease onset (Suppl. Figure-5). However, this observation was made in neuronal SH-SY5Y cells and incomparable to human or rat physiological / pathological conditions. Therefore, it needs further investigation in more relevant genetically modified or AAVs-induced disease models. We propose that the observed increase in level of ubiquitin during diseased conditions may be transitory and time dependent study would shed light in this direction. Altogether, UBA52 overexpression in neuronal cells prevents the lapse in proteasome activity during PD onset therefore, maintaining the protein turn-over and inhibiting protein aggregation.

### Effect of UBA52 on client proteins of HSP90 and ER stress related pathophysiological state

Given the powerful ability of Myc-UBA52 in protecting SH-SY5Y cells against altered expression of PD specific pathological markers, we further checked the effect of UBA52 on the protein level of HSP90 in SH-SY5Y cells. Myc-UBA52 significantly reduced the augmented protein level of HSP90 and this reduction was more prominent in rotenone-induced Myc-UBA52 group and maybe in order to prohibit the neurodegeneration (Fig 8a). This finding is in agreement to a previous report by Alam et al. 2017) suggesting that HSP90 inhibition may be a potential target in treating diverse array of neurodegenerative diseases.

**FIGURE 8:**
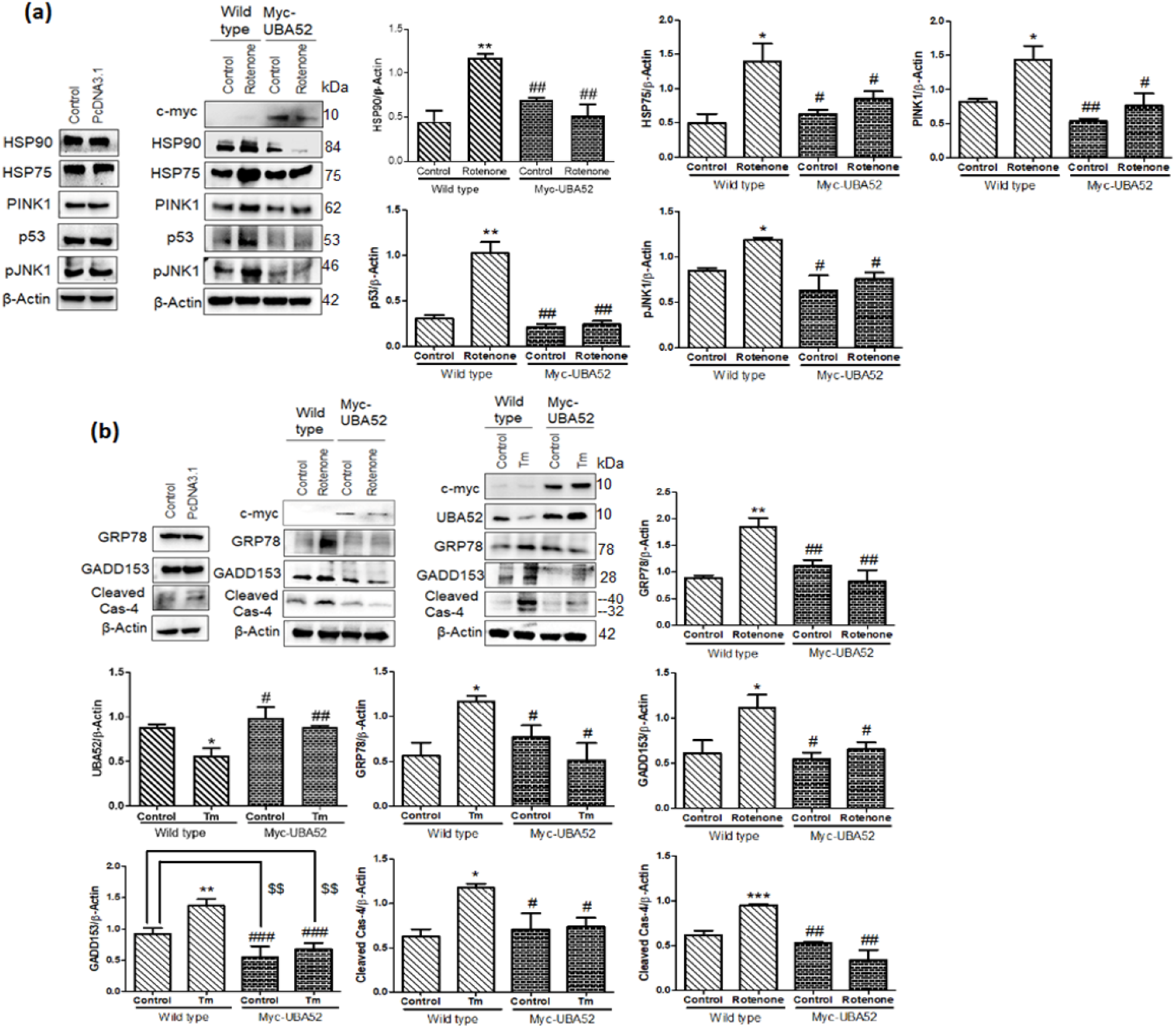
HSP90 client proteins and ERS (ER stress) marker proteins expression in UBA52 overexpressed SH-SY5Y cells. **(a)** SH-SY5Y cells were transfected with PcDNA3.1 or Myc-UBA52 with or without rotenone treatment and protein expression of client proteins of HSP90 were immunoblotted with indicated antibodies: n_exp_= 3, (b) Blots represent the protein expression of ER stress markers in PcDNA3.1 or Myc-UBA52 transfected SH-SY5Y cells with or without rotenone or tunicamycin (Tm) treatment. Graphs indicate the statistical analysis of various proteins normalized against β-Actin; n_exp_=3. Quantification are mean and SEM of at least three independent experiments and statistical analysis were performed using two way-ANOVA, followed by Tukey’s multiple comparison test. * p<0.05, ** p<0.01, *** p<0.001 control vs. Rotenone; # p<0.05. ## p<0.01, ### p<0.001 Myc-UBA52 vs. Rotenone; $$ p<0.01 control vs. Myc-UBA52.

HSP90 associates with and stabilize more than 200 client proteins such as p53 (cellcycle), PINK1, AKT1, JNK1 and others whose expression is altered during PD pathogenesis (Park et al. 2011; Taipale et al. 2012; Pratt et al. 2014). Also, proteins such as HSP75/TRAP1 (isoform of HSP90 in mitochondria) and PINK1 participate in mitochondrial function and bioenergetics which severely compromised during PD (Shin & Oh 2014). In this context, we have observed the increased level of PINK1, p53, pJNK1 and HSP75/TRAP1 in neuronal cells upon neurotoxin exposure which was inhibited in Myc-UBA52 transfected cells even after PD induction, suggesting the involution of UBA52 in HSP90 and its client proteins interaction during PD onset (Fig 8a). In view of significant investigation of ER stress signaling in PD pathogenesis (Goswami et al. 2016; Gupta, Mishra, et al. 2021; Singh et al. 2021; Ryu et al. 2002; Holtz & O’Malley 2003), further study was expanded to assess the effect of UBA52 on ER stress-related signaling factors such as GRP78, GADD153, cleaved caspase-4 (human analogue of caspase-12) in control and Myc-UBA52 transfected cells upon PD induction. Transient expression of Myc-UBA52 in neuronal cells inhibited the PD-related upregulated level of ER stress markers (Fig 8b). Moreover, as a positive control, we performed similar set of experiment using tunicamycin (ER stress inducer) to study the possible role of UBA52 in ER stress related signaling. As shown in Fig 8b, no evident difference was observed in Myc-UBA52 transfected neuronal cells upon tunicamycin treatment compared to only tunicamycin-treated neuronal cells. Findings indicated that UBA52 overexpression protects the neuronal cells against UPS initiation and chaperone related PQC during diseased condition and may have therapeutic implications.

## Discussion

Considering protein misfolding as a prime pathology in PD pathogenesis, the available information regarding UPS and its regulatory components (ubiquitin and ubiquitin ligases) is limited. Ubiquitin ligases are essential in mandatory tagging of ubiquitin to target proteins for their processing to degradation via proteasome. To date, large number of ubiquitin ligases have been reported but their known participation in PD pathology is scarce. In PD pathology, the important role of Parkin, CHIP, E6AP, SIAH, ITCH, TRIM2/9 has been reported (Tiwari & Singh 2020; Shin et al. 2005; Mund et al. 2018) and that to in confined test systems. Although, the function of E3 ligase Parkin is extensively studied in association with PINK1 to regulate the energy biogenesis (Rohé et al. 2004; Wauer et al. 2015; Truban et al. 2017), however, the role of the core component of UPS that is ubiquitin has not been yet reported. In this study, we for the first-time report the significant role of UBA52 in PD pathology during the early phase of disease. Additionally, we divulge the participation of UBA52 in CHIP-mediated ubiquitylation of molecular chaperone, HSP90 and finally decipher the role of UBA52 in orchestrating the associated neurodegenerative signaling. Separately, we showed that transient overexpression of UBA52 in neuronal cells provides protection against onset of PD pathology, highlighting its critical partaking in initiation of death mechanism probably due to protein misfolding and insufficient chaperoning, eventually protecting the dopaminergic neurons.

Our findings in human Myc-α-synuclein transfected neurons, α-synuclein-PFFs treated SHSY5Y cells, rotenone-induced sporadic cellular and rat models of PD and SNCA transgenic mice highlights that UBA52 is downregulated during early phase of sporadic PD which is critical and might account for the reduction in proteasome activity during the diseased condition. UBA52 on one hand, interacts with an important PD-related pathological protein, α-synuclein and inhibits its upregulation to regulate its level to impede the fibril formation. On the other hand, overexpression of UBA52 in both proliferative dopaminergic SH-SY5Y cells and differentiated SH-SY5Y, N2a and PC12 neuronal cells averts the downregulation of TH and caspase-3 activation. These findings raise the possibility that UBA52 has a perspicacious role in early progression of PD related pathogenesis. Our previous study on the role of ER stress (Goswami et al. 2016; Gupta, Mishra, et al. 2021) and on reduced proteasome activity in PD pathology (Singh et al. 2021) advocated the probable role of molecular chaperone, as both ER functioning and PQC are closely regulated by chaperones in neuronal cells (Bhattacharyya et al. 2014). Among several chaperones, HSP90 has emerged as a novel therapeutic target for PD treatment (Alam et al. 2017). In concurrence to this, our quantitative mass spectrometric and co-immunoprecipitation data demonstrated that UBA52 interacts with HSP90 and CHIP, forming a complex, wherein we suggested that E3 ligase CHIP adds K63-mediated ubiquitin chain to HSP90 in the presence of UBA52. K63-mediated ubiquitylation is reported to be associated with various post-translational modifications like endocytosis, signal transduction and autophagy (Erpapazoglou et al. 2014). It has been previously reported that K63-mediated ubiquitylation also reduces the cargo stress from proteasome and facilitates the clearance of inclusion bodies and supports the mitochondrial homeostasis (Lim & Lim 2011) and endorse our findings in context to observed UBA52 overexpression-mediated neuroprotection. Despite the downregulation of UBA52 level during acute phase of disease onset, we propose that CHIP-UBA52 allied K63-linked HSP90 ubiquitylation spikes during sporadic PD conditions due to increase in protein misfolding, HSP90 protein level and to maintain the homeostatic protein turnover, highlighting the indispensable role of UBA52 in maintaining HSP90 level. However, progression of disease and initiation of other disease-related neurodegenerative signaling mechanisms tardily leads to cell death and needs to be investigated in detail. In agreement to our previous finding, a recent study has shown that the proteasome hydrolytic activity reduces during PD pathology, acting either as cause or consequence to further exacerbate the disease pathology (Bi et al. 2021; Singh et al. 2021). Protection against reduced proteasomal activity after rotenone exposure in the presence of overexpressed UBA52 also reflects its burgeoning part in sufficient degradation of target proteins such as HSP90 and impede neurodegeneration during early PD. Our most surprising study is that control neuronal cells in which UBA52 was transiently silenced had the reduced total ubiquitin amount. However, neuronal cells expressing disease markers after exposure to the neurotoxin compensated for the loss of UBA52 with a high level of total ubiquitin content. In view of this unexpected finding, we assume that onset of disease increases the cargo load on the proteosome which boost the total ubiquitin level to temporarily escalate the protein turnover in order to restore proteostasis, compensating for the loss of UBA52. Although, our study was conducted in acute phase of diseased conditions, we observed that the proteasome activity was downregulated analogous to upregulated ubiquitin levels during disease onset. Therefore, a detailed time dependent study may shed light in this context. Since UBA52 depletion is observed at the early phase of disease pathogenesis, the diagnostic aspect of UBA52 could also be explored though it may need excellent technical tools and specific dyes. However, it is an apparently achievable target in account of an investigation regarding its peripheral level in the blood, urine and cerebrospinal fluid. Recently, (Xu et al. 2021) have also suggested the clinical significance of UBA52 level in urine samples for the diagnosis of diabetes mellitus and diabetic nephropathy.

Our findings related to augmented HSP90 protein abundance during PD are in coherence to previous findings underlining that HSP90 inhibitor, Geldanamycin (GA) potentially reduces the α-synuclein oligomer and aggregate formation as well as induces α-synuclein clearance (Ebrahimi-Fakhari et al. 2011). Similarly, the other HSP90 inhibitor 17-AAG (17-allylamino-17-demethoxygeldanamycin, or Tanespimycin) also reduces the α-synuclein toxicity and facilitates its clearance in various experimental models of PD (Ebrahimi-Fakhari et al. 2011). Consistent with these research findings and our data that UBA52 surplus in neuronal cells resists the upregulated level of HSP90, we proffer UBA52 as a promising candidate for addressing PD and Parkinsonism-related disorder. Also, our findings affirm the previous findings on HSP90 inhibition, suggesting that HSP90 co-localize with α-synuclein and UBA52 surplus forestalls the protein aggregation by interacting with both the aforementioned proteins and regulating their levels.

To broaden the research arena, we also evaluated the level of few bona fide client proteins of HSP90 which participate in PD-associated various neurodegenerative signaling such as pJNK1, p53, PINK1 (PARK6). Myc-UBA52 transfected neuronal cells inhibited the change in the protein level of studied client proteins, suggesting that UBA52 has a salient role in pathways related to cell cycle arrest, apoptosis, autophagy and mitochondrial homeostasis. Mitochondrial chaperone, HSP75/TRAP1 regulates the formation of reactive oxygen species (ROS) and shows explicit connection with PINK1, wherein PINK1 phosphorylates HSP75 to protect the neuronal cells against oxidative stress (Pridgeon et al. 2007). Findings showed insignificant alteration in the protein level of HSP75 after exposure to rotenone in Myc-UBA52 transfected cells. Altogether, our data points that UBA52 implicitly or explicitly regulates the mitochondrial functioning and has a multi-facet role in maintaining cellular machinery to prevent the onset of disease. Our confocal microscopy-based observations also showed localization of UBA52 in the mitochondrion during disease onset, reaffirming the implications of UBA52 in mitochondrial homeostasis. Since protein homeostasis is essential for cellular structure and function therefore, protein misfolding or unfolding (as observed in PD pathology and other neurodegenerative diseases) intends to activate the cellular PQC involving various molecular chaperones and UPS (Bhattacharya et al. 2020). It is evident from our generated data in cellular and rodent PD model that UBA52 surfeit reduces the proteotoxic stress and regulate the ER functionality, as evident by the insignificant change in the protein level of ER stress markers upon exposure to the neurotoxin to prevent the neuronal death.

In summary, our findings propose conspicuous role of UBA52 in maintaining the E3 ubiquitin ligase CHIP-mediated ubiquitylation of chaperone HSP90, along with subsequent effect on its important client proteins, thereby regulating multiple pathways simultaneously during PD pathogenesis (Fig 9). Such diverse role of UBA52 should be explored further in the context to its diagnostic, pharmacological and translational aspects. Though the study was focused on human Myc-α-synuclein/UBA52 transfected neurons, α-synuclein-PFFs and rotenone-induced cells and rat models of PD and SNCA transgenic mice, the limitation of the current study is non-availability of UBA52 specific transgenic animal model to assess its effect on PD pathology especially intracellular signaling pathways and mitophagy/autophagy which are very challenging and subject to future investigation. Nevertheless, our study claims that UBA52 displays significant efficacy in orchestrating various signaling mechanisms related to dopaminergic neuronal death, in view of its direct interference in regulating the α-synuclein level and ubiquitylation of HSP90 in complex with CHIP to maintain the efficient chaperoning. In addition, other neurodegenerative diseases in which the therapeutic implications of HSP90 inhibitors have been suggested may perhaps consider UBA52 as a promising candidate for targeting disease therapeutics and mitigating neuronal death.

## Abbreviations

PD: Parkinson’s Disease
SN: Substantia nigra
STR: Striatum
TH: Tyrosine hydroxylase
UPS: Ubiquitin Proteasome System
ERS: Endoplasmic Reticulum Stress
UBA52: Ubiquitin-60S ribosomal protein L40
UBB: Polyubiquitin-B
UBC: Polyubiquitin-C
RPS27a: Ubiquitin-40S ribosomal protein S27a
RT-qPCR: Real time quantitative PCR
MS: Mass Spectrometry
MALDI-TOF: Matrix assisted laser desorption ionization-Time of flight
ACN: Acetonitrile
ABC: Ammonium Bicarbonate
TFA: Trifluro acetic acid
IgG: Immunoglobulin G
PQC: Protein Quality Control
PSOPIA: Prediction server of protein-protein interaction
HSP90: Heat Shock Protein 90
CHIP: C-terminus of HSC-70
K^48^R: Lysine residue mutated to Arginine at position-48 of UBA52
K^63^R: Lysine residue mutated to Arginine at position-63 of UBA52
pJNK1: phospho-c-Jun N-terminal kinase 1
AKT1: RAC-alpha serine/threonine-protein kinase 1
PINK1: PTEN-induced kinase 1
HSP75: Heat Shock Protein 75

## Affiliations

^1^Division of Neuroscience and Ageing Biology, Division of Toxicology and Experimental Medicine, CSIR-Central Drug Research Institute, Lucknow 226031, India ^2^Academy of Scientific & Innovative Research (AcSIR), Ghaziabad 201002, India.

**Correspondence Author**

Dr Sarika Singh

## Author Contribution

Shubhangini Tiwari and Abhishek Singh conceptualized, designed the work plan and performed the experiments; Parul Gupta contributed in the experimental work and Shubhangini Tiwari wrote the manuscript; Sarika Singh conceptualized, designed and supervised the study as well as reviewed and edited the final manuscript.

## Acknowledgment

We thank Dr Ramesh Sharma, Ms Anupma Saxena, Ms Kavita Singh, Mr CP Pandey and Mr Toofan Raut and SAIF facility (CSIR-CDRI) for technical assistance during experiments. We also acknowledge University Grants Commission (UGC), New Delhi, India for providing one-time research grant and Jawahar Lal Nehru University for providing the research opportunity.

## Ethics approval

Non-transgenic SD rats were used after approval from the Institutional animal ethics committee of CSIR – Central Drug Research Institute (IAEC/2018/F-52), whereas, SNCA C57BL/6J-Tg (Th-SNCA*A30P*A53T) 39Eric/J transgenic mice (Stock number: 008239) were purchased from the Centre of Cellular and Molecular Biology (CCMB), Hyderabad [CSIR-CDRI (IAEC/2021/SI no2)].

*For humans: Not applicable.

## Consent for publication

All authors read and approved the final manuscript.

## Availability of data and material

All the generated data is compiled and given in MS or Supplementary file. The raw data will be provided on request.

---

**Figure.**
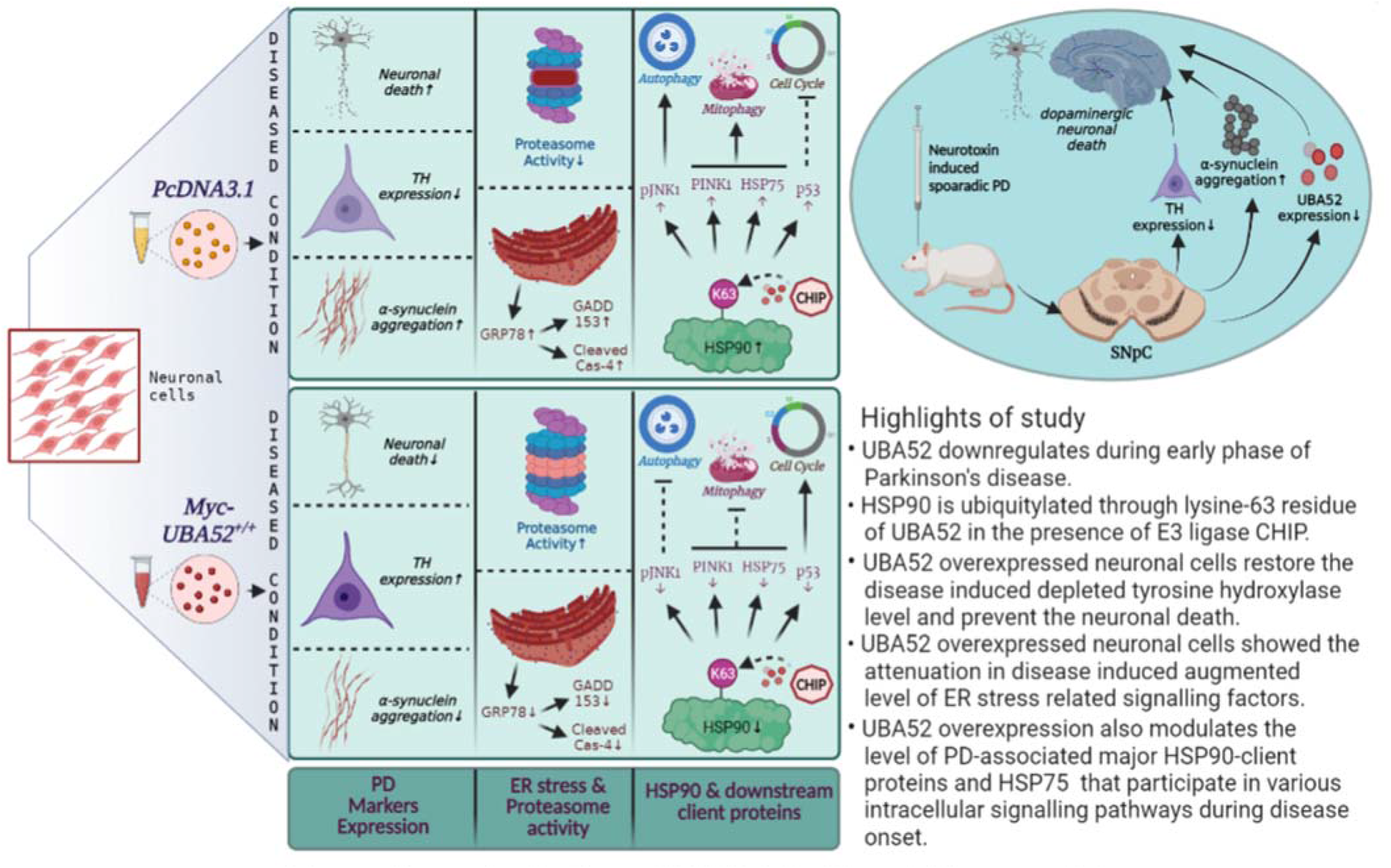
Diverse implications of UBA52 in Parkinson’s Disease pathology

