## Supplementary Data for "UBA52 is crucial in HSP90 ubiquitylation and neurodegenerative signaling during early phase of Parkinson’s disease"

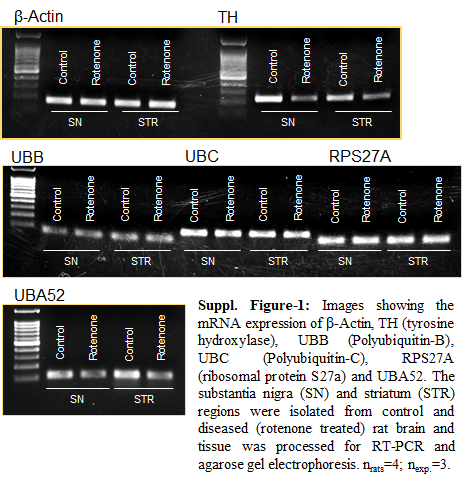

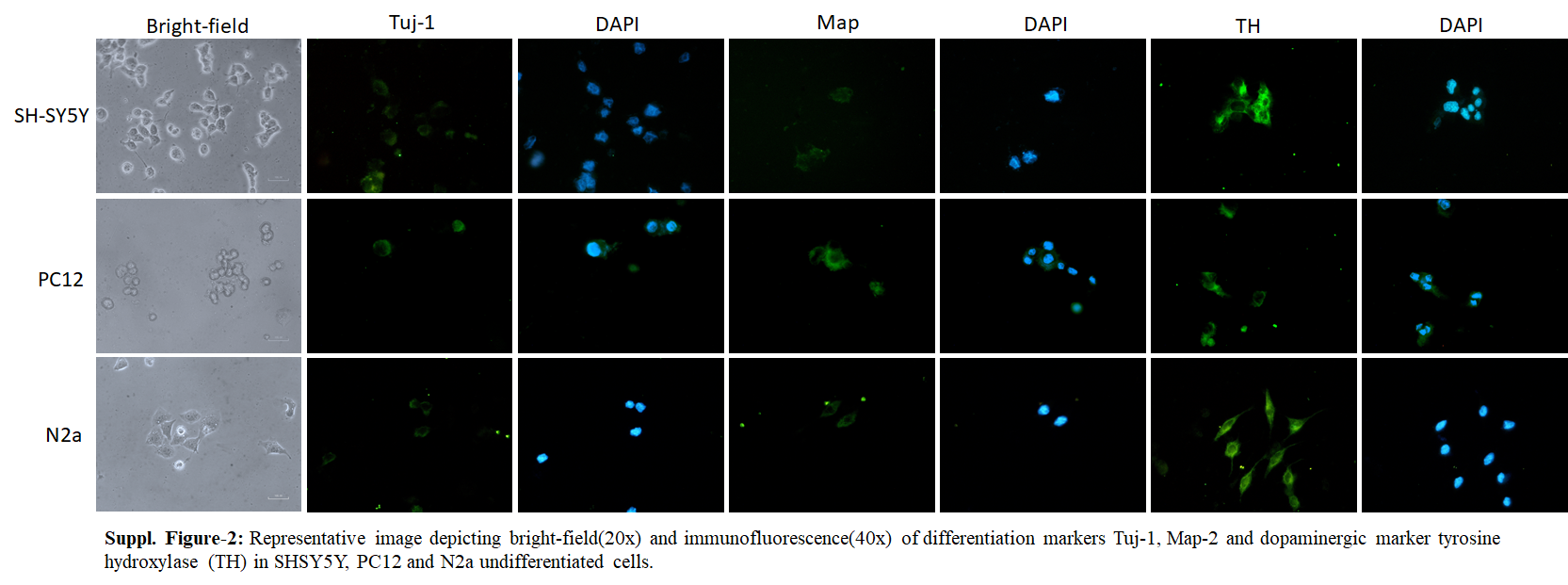

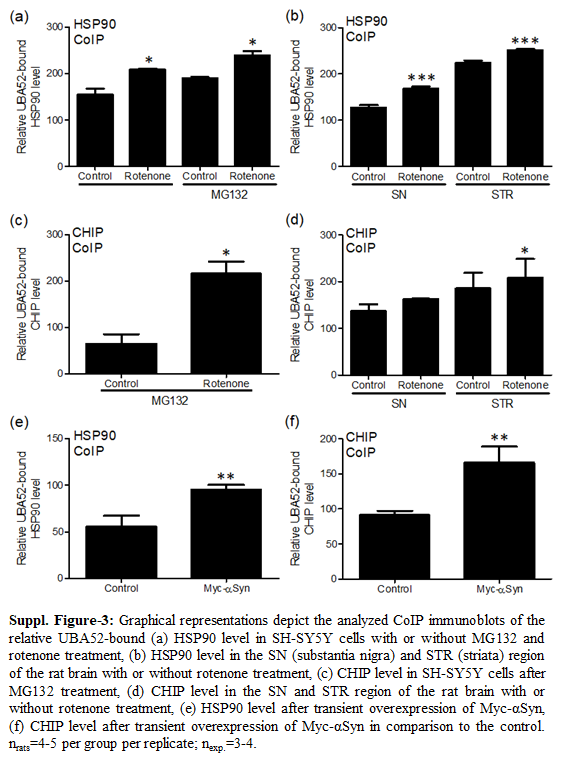

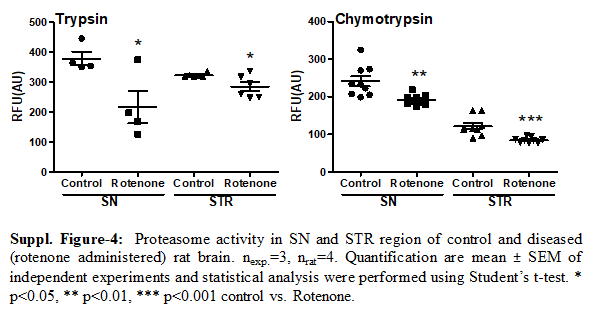

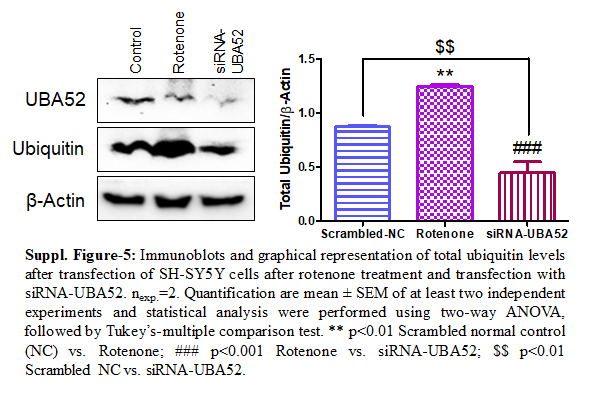

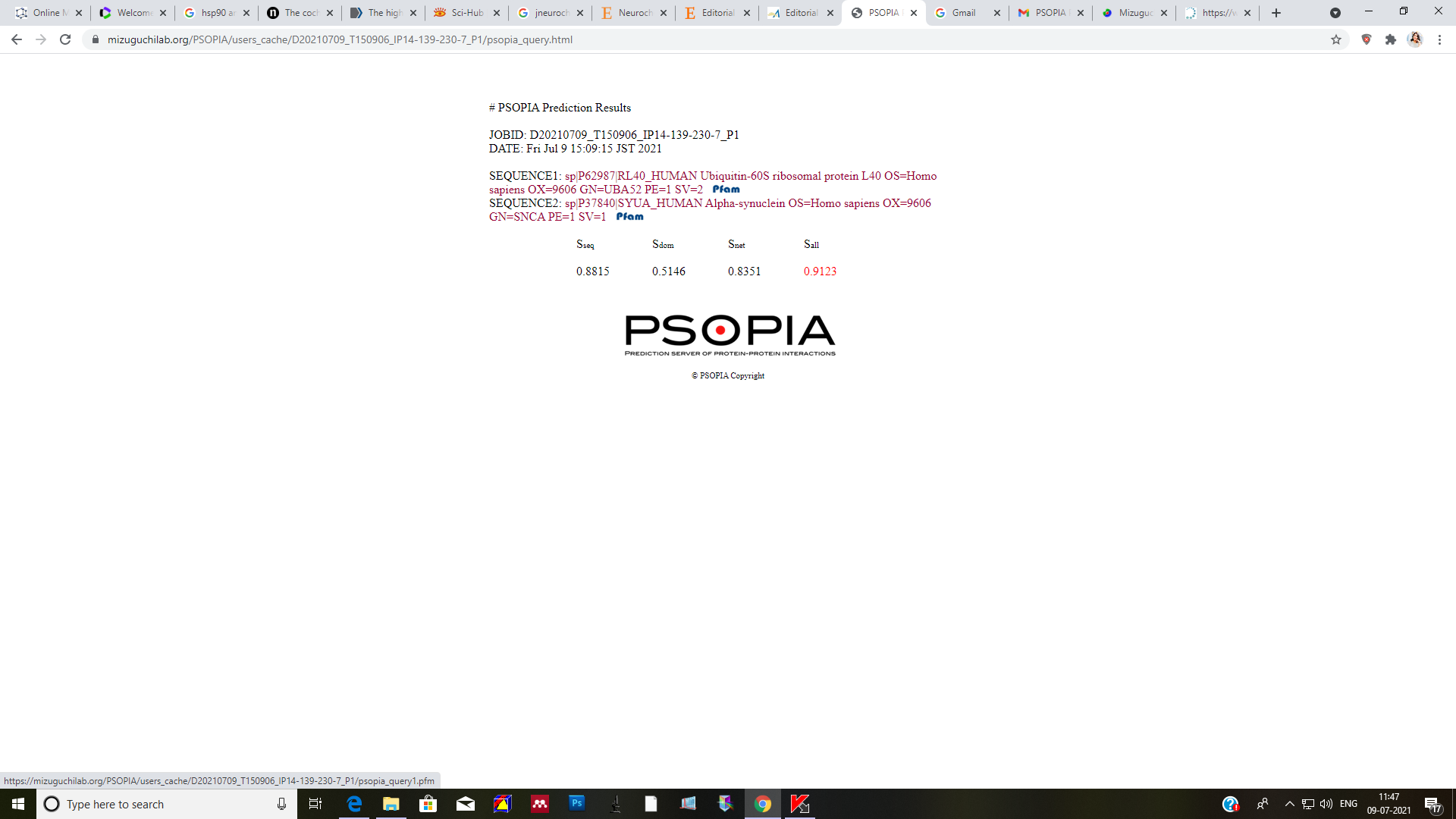

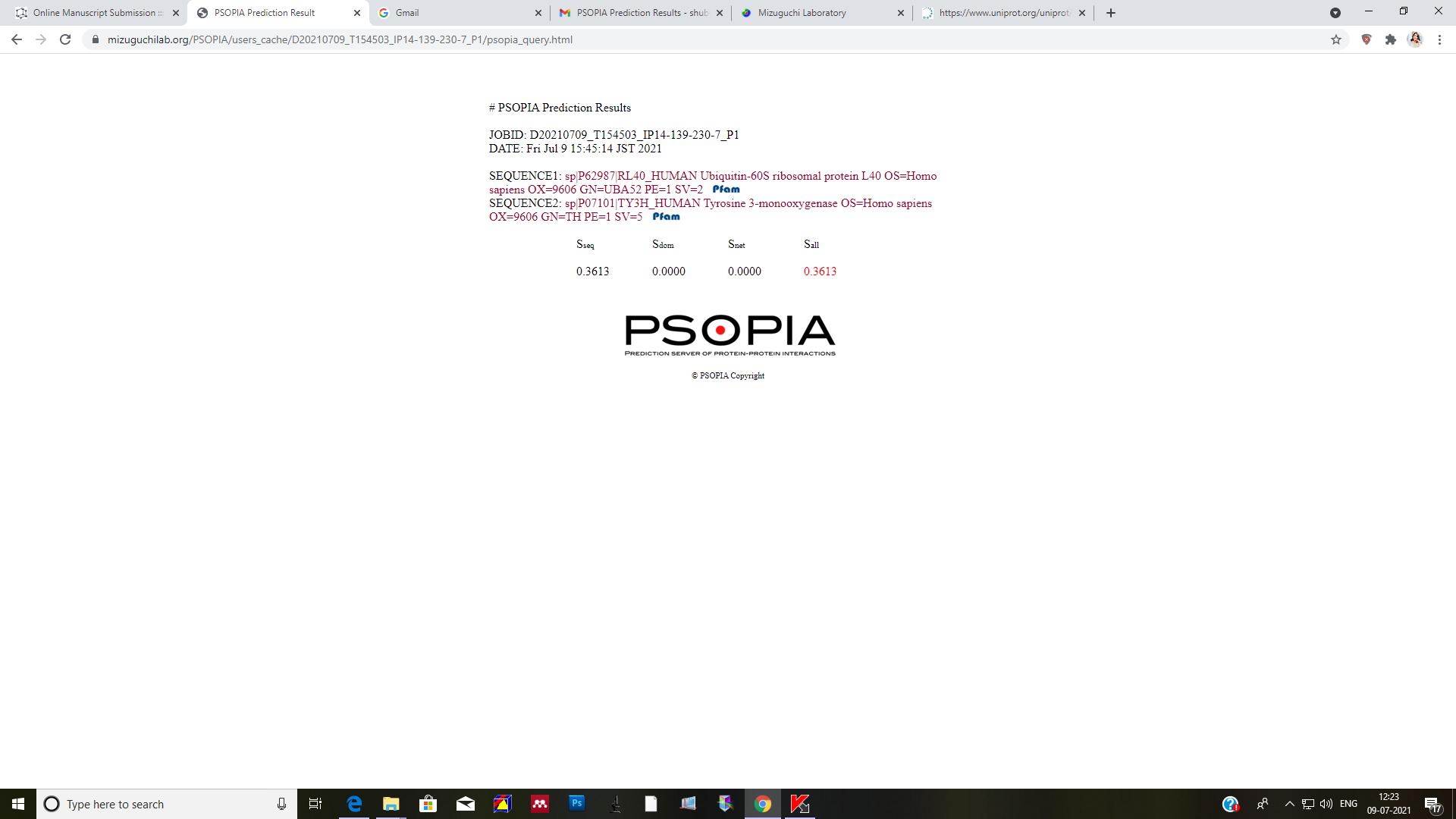

a.

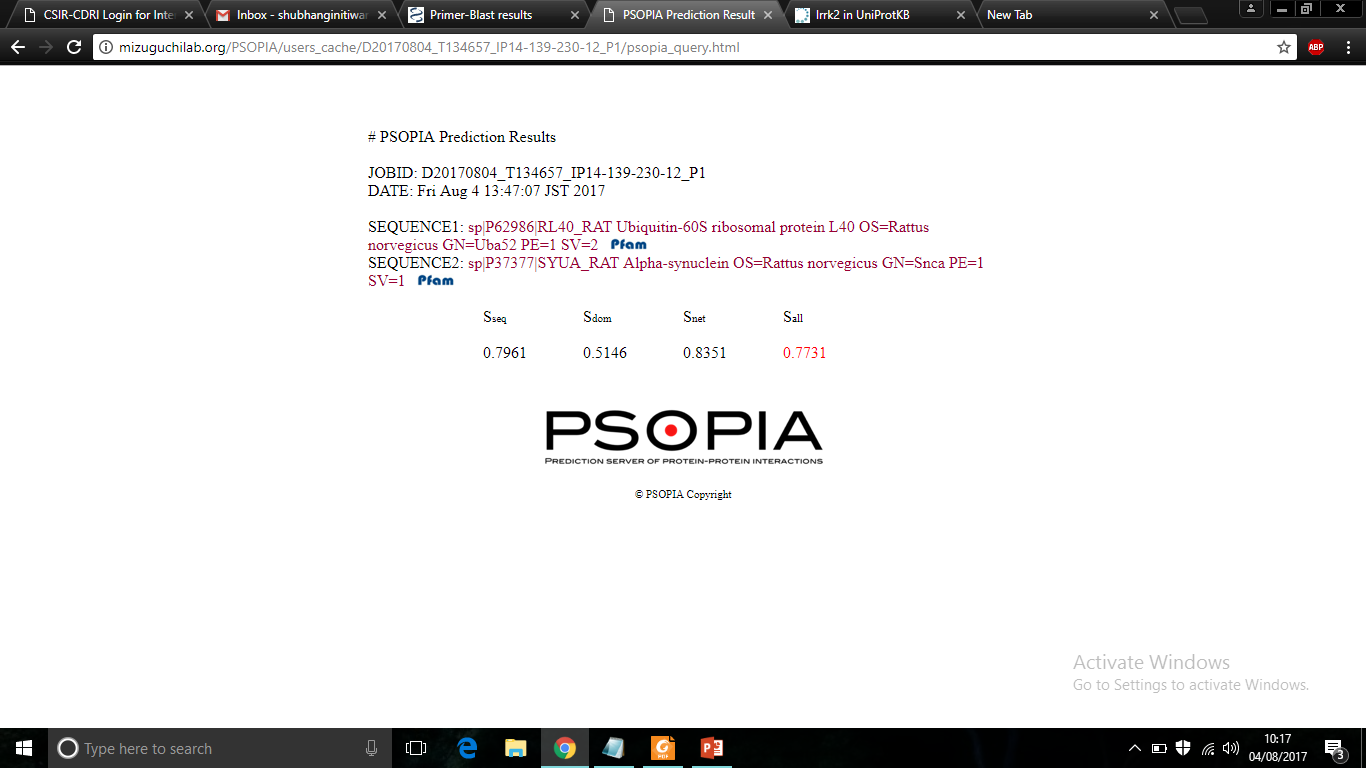

b.

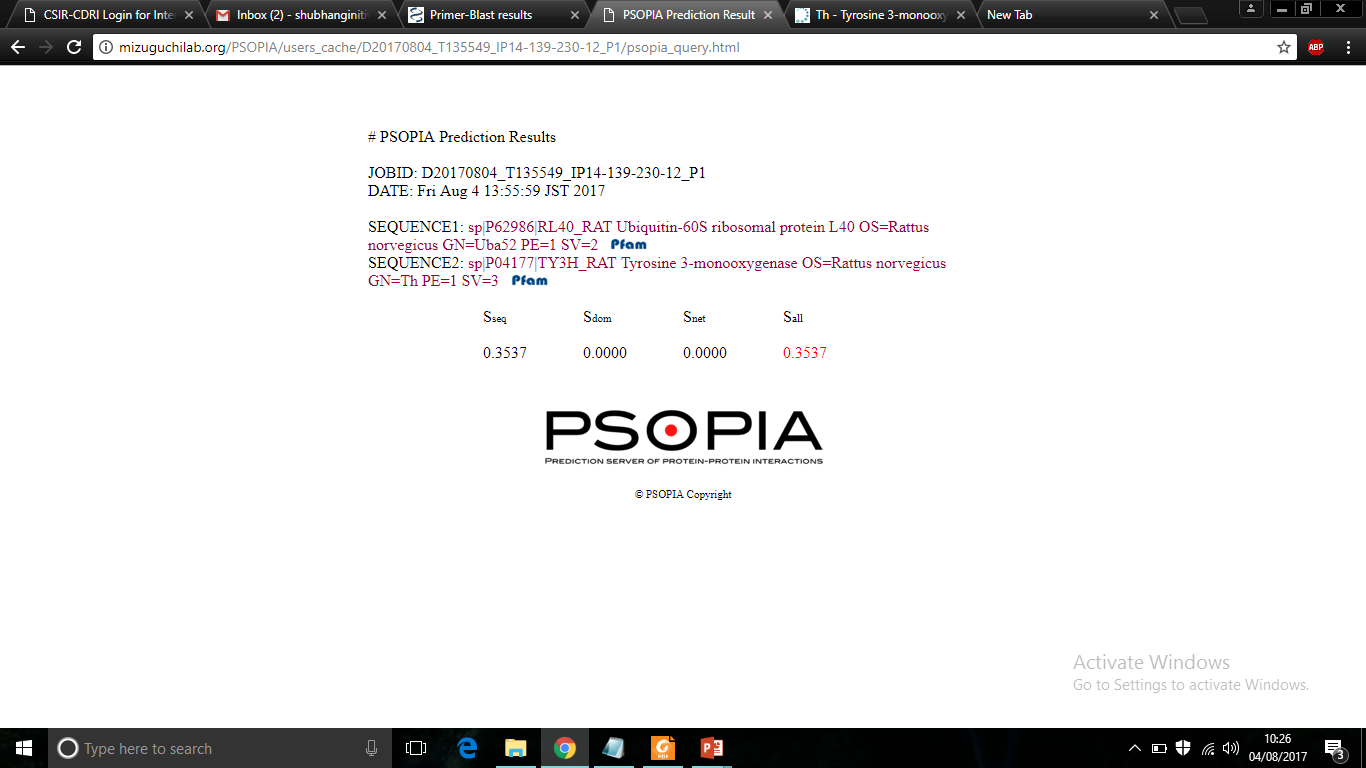

**Suppl. File-1:** In-silico analysis indicating the interaction of UBA52 with TH and α-synuclein as assessed through a freely accessible server, PSOPIA (Prediction Server of Protein-Protein Interaction), based on both human **(a)** and rat **(b)** protein sequences available on UniProtKB database.

**Supplementary Information:**

Chemicals:

Bovine serum albumin (BSA), disodium hydrogen phosphate (Na_2_HPO_4_), dimethyl sulfoxide (DMSO), ethidium bromide, glucose,4-(2-hydroxyethyl)-1-piperazine ethane sulfonic acid (HEPES), 3-(4,5-dimethylthiazol-2-yl)-2,5- diphenyltetrazolium bromide dye (MTT), NP-40, phenylmethylsulphonyl fluoride (PMSF), magnesium chloride (MgCl_2_), dithiothreitol (DTT), RNase, sodium bicarbonate and tris-buffer were procured from SRL, India. Dulbecco's modified Eagle's medium (DMEM), fetal bovine serum (FBS), Trizol, Ham's F12 medium, penicillin-streptomycin, nuclease-free water, lipofectamine 3000 and mitotracker-red and -deep red were purchased from Invitrogen (San Diego, CA, USA). Luria Bertani broth and agar powder were purchased from Himedia. *In vitro* ubiquitylation kit was purchased from Enzo-Life sciences. Recombinant human α-synuclein protein PFF (ab218819) and recombinant mouse α-synuclein protein PFF (ab246002) were purchased from Abcam. Other chemicals such as Protein-A Sepharose beads, anti-fade medium DAPI, copper sulfate (CuSO_4_), calcium chloride (CaCl_2_), Folin–Ciocalteu reagent, potassium chloride (KCl), sodium carbonate (NaHCO_3_), sodium chloride (NaCl), sodium dihydrogen phosphate (NaH_2_PO_4_), sodium hydroxide (NaOH), protease and phosphatase inhibitor cocktail, rotenone, tunicamycin, apomorphine, paraformaldehyde (PFA), ethylenediaminetetraacetic acid (EDTA), acetonitrile (ACN), ammonium bicarbonate (ABC), trifluoro acetic acid (TFA) and trypsin MS grade were obtained from Sigma, USA.

Table 1: Antibodies used for Immunoblot (WB) and immunofluorescence (IF)

| Antibodies | Company | Catalogue  Number | Species | Dilution  WB | Dilution  IF |
| --- | --- | --- | --- | --- | --- |
| β-Actin | Sigma-Aldrich | A3854 | Mouse | 1:10000 |  |
| α-Synuclein | Santa Cruz  Biotechnology | sc-53955 | Mouse | 1:500 | 1:100 |
| TH | Sigma-Aldrich | T8700 | Rabbit | 1:1000 |  |
| TH | Sigma-Aldrich | T2928 | Mouse | 1:1000 | 1:250 |
| UBA52 | Abcam | ab109227 | Rabbit | 1:1000 | 1:250 |
| HSP90 | Cell Signalling | 4874S | Rabbit | 1:1000 |  |
| HSP90 | Santa Cruz  Biotechnology | sc-515081 | Mouse | 1:500 | 1:100 |
| CHIP | Santa Cruz  Biotechnology | sc-133066 | Mouse | 1:500 |  |
| pJNK | Santa Cruz  Biotechnology | sc-6254 | Mouse | 1:500 |  |
| p53 | Santa Cruz  Biotechnology | sc-98 | Mouse | 1:500 |  |
| Cleaved Caspase-3 | Invitrogen | 700182 | Rabbit | 1:1000 |  |
| Cleaved  Caspase-4 | Santa Cruz  Biotechnology | sc-56056 | Mouse | 1:500 |  |
| Ubiquitin | Cell Signalling | 3936S | Mouse | 1:1000 |  |
| Myc-Tag | Cell Signalling | 2276S | Mouse | 1:1000 |  |
| Flag-M2 | Sigma | F1804 | Mouse | 1:1000 |  |
| HSP75 | Santa Cruz  Biotechnology | sc-13577 | Rabbit | 1:1000 |  |
| PINK1 | Santa Cruz  Biotechnology | sc-517353 | Mouse | 1:500 |  |
| GRP78 | Abcam | ab21685 | Rabbit | 1:5000 |  |
| GADD153 | Santa Cruz  Biotechnology | sc-7351 | Mouse | 1:500 |  |
| Anti-Mouse Secondary | Sigma-Aldrich | A9044 |  | 1:5000 |  |
| Anti-Rabbit Secondary | Sigma-Aldrich | A0545 |  | 1:3000 |  |
| Alexa-fluor 488 Green | Invitrogen | A11034 | Rabbit |  | 1:300 |
| Alexa-fluor  488 Green | Invitrogen | A11059 | Mouse |  | 1:300 |
| Alexa-fluor 546 Red | Invitrogen | A11003 | Mouse |  | 1:300 |
| Alexa-fluor 647 Deep Red | Invitrogen | A21235 | Mouse |  | 1:300 |

Table 2: List of primers sequences used for RT-PCR

| Gene Name | cDNA Primer Sequences |
| --- | --- |
| β-actin | 5' GTCGTACCACTGGCATTGTG 3'  3' CTCTCAGCTGTGGTGGTGAA 5' |
| TH (Human) | 5' TGTGGCCTTTGAGGAGAAGGA 3'  3' TCAAACACCTTCACAGCTCGG 5' |
| TH (Rat) | 5' CCA CGG TGT ACT GGT TCA CT 3'  3' GGC ATA GTT CCT GAG CTT GT 3' |
| UBA52 (Human) | 5' AGATGATGCCAAAGGACGCA 3'  3' TCAATGGTGTCACTGGGCTC 5' |
| UBA52 (Rat) | 5' GCAGACGCCAACATGCAGA 3'  3' GGGGATGCCTTCCTTGTCTT 5' |
| RPS27A (Rat) | 5' GGTCTAATCCGTCTCTTTTC 3'  3' CTGGATCTTGGCCTTTACAT 5' |
| UBB (Rat) | 5' TAGCCATTTGCCCCAATTTA 3'  3' TGCTTACCATGCAACAAAAC 5' |
| UBC (Rat) | 5' ACCTTTCTCACCACAGTATC 3'  3' AAACTAAGACACCTCCCCAT 5' |
| Tg-SNCA-F  Tg-SNCA-R  (Transgene: 469 bp  WT: No-bands) | 5′-CAG GTA CCG ACA GTT GTG TAA AGG AAT-3′  5′-GAT AGC TAT AAG GCT TCA GGT TCG TAG TCT-3′ |

Note: Confirmation of plasmid transformation was done using primers provided in the kit by OriGene technologies and plasmid transfection in SH-SY5Y cells was confirmed using primers stated above.
